# A haplotype-resolved chromosome-level assembly and annotation of European hazelnut (*C. avellana* cv. Jefferson) provides insight into mechanisms of eastern filbert blight resistance

**DOI:** 10.1101/2023.09.21.558899

**Authors:** S.C. Talbot, K.J. Vining, J.W. Snelling, J. Clevenger, S.A. Mehlenbacher

## Abstract

European hazelnut (*Corylus avellana* L.) is an important tree nut crop. Hazelnut production in North America is currently limited in scalability due to *Anisogramma anomala,* a fungal pathogen that causes Eastern Filbert Blight (EFB) disease in hazelnut. Successful deployment of EFB resistant cultivars has been limited to the state of Oregon, where the breeding program at Oregon State University (OSU) has released cultivars with a dominant allele at a single resistance locus identified by classical breeding, linkage mapping, and molecular markers. ‘Jefferson’ is resistant to the predominant EFB biotype in Oregon and has been selected by the OSU breeding program as a model for hazelnut genetic and genomic research. Here, we present a near complete, haplotype-resolved chromosome-level hazelnut genome assembly for *C. avellana* ‘Jefferson’. This new assembly is a significant improvement over a previously published genome draft. Analysis of genomic regions linked to EFB resistance and self-incompatibility confirmed haplotype splitting and identified new gene candidates that are essential for downstream molecular marker development, thereby facilitating breeding efforts.

## Introduction

European hazelnut (*Corylus avellana* L.) is an important specialty tree nut crop that is grown in temperate climates for use in the in-shell and kernel markets, typically consumed raw or roasted, in confectionaries and baked goods. The estimated value of the global hazelnut industry is three billion US dollars with Turkey representing nearly 70% of global production (FAO, 2022). Hazelnut (2n = 2x = 11) is a woody perennial that is monecious, dichogamous, wind-pollinated, and self-incompatible (Hill et al., 2021). While all hazelnut species produce edible nuts, the European hazelnut (*Corylus avellana* L.) is the most widely grown because of its desirable characteristics such as a large high-quality nuts, thin shells, and desired flavor profile. Traditional cultivars are clonally propagated and originated as selections from the wild in Europe and western Asia (Mehlenbacher and Molnar, 2021).

Commercial hazelnut production in North America has been limited due to the high susceptibility of European hazelnut to *Anisogramma anomala,* a biotrophic ascomycete, and the causal agent of the eastern filbert blight (EFB) disease. *A. anomala* has co-evolved with its endemic host, the American hazelnut (*Corylus americana*), and in the wild, the disease is widely tolerated (Capik and Molnar, 2012; Revord et al., 2020). Symptoms of EFB are apparent ∼18 months following initial infection, and include branch die-back, girdling of trunks, and eventual tree and orchard death. While management techniques such as pruning, scouting for cankers, and applying fungicides can slow the disease’s spread, they do not eliminate it (Pscheidt and Ocamb, 2022). Thus, breeding for genetic resistance is considered the most sustainable approach to managing EFB.

Oregon State University (OSU) has been a leader in developing improved EFB resistant cultivars for the Pacific Northwest (PNW), where Oregon represents 95% of US hazelnut production. The OSU hazelnut breeding program’s primary contribution to EFB-resistant cultivar development can be traced to a 1975 discovery in southwest Washington of the obsolete pollinizer, ‘Gasaway’, which was completely free of EFB in a highly infected and dying orchard of ‘DuChilly’ (Thompson et al., 1996). To date multiple resistant pollinizers and cultivars derived from ‘Gasaway’ have been released (Mehlenbacher, 2021), and underlie the expansion of acreage planted in Oregon, which increased from ∼11,000 ha in 2009 to greater than 25,000 ha in 2022 (USDA-NASS, 2023). Outside of Oregon, however, cultivars with ‘Gasaway’ resistance have been shown to be susceptible to genetically diverse *A. anomala* populations (Muehlbauer et al., 2019). Indeed, a genome assembly of the pathogen has shown that it has one of the largest Ascomycota genomes suggesting a high capacity for pathogenic variation (Cai et al., 2013). The long-term durability of Oregon’s commercial hazelnut orchards and the potential for expanding hazelnut production is limited by the pathogen’s variability and narrow resistance offered by ‘Gasaway’.

The availability of genomic resources in *Corylus* has been increasing in recent years. The cultivar ‘Jefferson’ was chosen for the first *Corylus* genome assembly because it contains ‘Gasaway’ EFB resistance and it was selected from the reference mapping population (Mehlenbacher et al., 2006). However, the Illumina-based first draft was highly fragmented due to hazelnut’s highly heterozygous nature and the limitations imparted by short-read sequencing and assembly technologies (Rowley et al., 2018). With advances in long-read sequencing, pseudo-chromosome level genome assemblies for *Corylus* have been made available for *C. avellana* cultivars ‘Tombul’ and ‘Tonda Gentile delle Langhe’ (Lucas et al., 2021; Pavese et al., 2021) and representative accessions of two *Corylus* species, *C. heterophylla* Fisch (Liu et al., 2021; Zhao et al., 2021) and *C. mandshurica* Maxim (Li et al., 2021). However, these genome assemblies are collapsed and there has been no haplotype-resolved “phased” assembly that represents both homologous chromosomes. Distinguishing between the two chromosomes is essential for determining the parental allelic contributions to self-incompatibility, EFB resistance, and other traits.

EFB resistance derived from ‘Gasaway’ has been characterized as a dominant allele at a single locus with 1:1 segregation (Mehlenbacher et al., 1991, 2006). This source of resistance has been mapped to linkage group (LG) 6 of the genetic map using random amplified polymorphic DNA (RAPD) and simple sequence repeat (SSR) markers in a segregating population from a cross between two heterozygous clones, susceptible ‘OSU 252.146’ x resistant ‘OSU 414.062’ (Mehlenbacher et al., 2006). From this mapping population, the elite cultivar ‘Jefferson’ was identified for release and was the source of the first *Corylus* draft genome (Mehlenbacher et al., 2011; Rowley et al., 2012). Fine mapping of the ‘Gasaway’ region using bacterial artificial chromosomes (BACs) identified a span of approximately 135 kb and five candidate EFB resistance genes (Sathuvalli et al., 2017). Other sources of EFB resistance have been identified and mapped in over 30 *C. avellana* cultivars and accessions, and while the majority map to LG6 (Sathuvalli et al., 2012; Colburn et al., 2015; Komaei Koma 2020), other sources of qualitative and quantitative resistance have been mapped to LG2 (Sathuvalli et al., 2011a; Şekerli et al., 2021), LG7(Bhattarai et al., 2017; Sathuvalli et al., 2011b; Şekerli et al., 2021), LG10 and LG11 (Lombardoni et al., 2022), and more recently LG4 and LG1 (unpublished). A complete summary of resistant cultivars and their related linkage group can be found in Table 1 of Mehlenbacher et al. (2023). The development of elite EFB resistant cultivars is a major goal in hazelnut breeding; however, the lengthy field evaluations provide more robust data on phenotypic variation. The accurate identification of candidate genetic parental contributions underlying qualitative and quantitative loci for EFB resistance will significantly aid in selection across a diverse collection of *Corylus* germplasm, thereby allowing for development of cultivars with multiple resistance loci.

**Table 1.** Summary statistics for the assembled *C. avellana* ‘Jefferson’ genomes.

| Statistics | 'Jeff V4 Hap1' | 'Jeff V4 Hap2' | "Jeff V4 Hap1"<br>Chr-resolved | "Jeff V4 Hap2"<br>Chr-resolved |
| --- | --- | --- | --- | --- |
| **Total Scaffold number** | 663 | 229 | 11 | 11 |
| **Total assembly Length (Mb)** | 385.8 | 372.5 | 349.7 | 352 |
| **N<sub>50</sub> (Mb)** | 23.4 | 22.5 | 32.5 | 32.4 |
| **Largest contig (Mb)** | 34.0 | 38.9 | 48.25 | 47.6 |
| **L50 (Mb)** | 7 | 7 | 5 | 5 |
| **Number of contigs merged** | NA | NA | 22 | 21 |
| **Number of predicted protein-coding genes** | NA | NA | 33,506 | 34,379 |

The largest class of characterized plant disease resistance (R) genes encode N-terminal Nucleotide Binding Site (NBS) and C-terminal Leucine-Rich-Repeat (LRR) functional domains (McHale et al., 2006). The LRR domain is highly variable within and among plant species and is typically associated with direct or indirect pathogen effector protein interactions (Prigozhin and Krasileva, 2021). R-genes are often localized into clusters within chromosomes and can have significant variations in encoded amino acid sequence motifs, even within specific categories of R-genes (Kroj et al., 2016; Bailey et al., 2018; Wang and Chai, 2020). Extensive research conducted over the past two decades has demonstrated the successful deployment of NBS-LRR R-genes in a wide range of crops (Kourelis and van der Hoorn, 2018). Investigating the complex molecular mechanisms of R-genes both within and across different plant species is an expensive and resource-intensive task. Past work has identified candidate EFB resistance genes in ‘Jefferson’, however, the functional descriptions are more than a decade old, and recent improvements in genome assembly, annotation algorithms, and curated databases of plant genomes represent an opportunity to improve the description of candidate R-genes. To better direct future research in the ‘Gasaway’ resistance region, it is crucial to update the annotation of *Corylus* R-gene candidates and evaluate them for protein domain similarities. This analysis will offer insights into the putative functionality of these genes and help determine if other sources of EFB resistance share similar molecular components.

Hazelnut orchard design and elite cultivar development also require an understanding of self-incompatibility. Hazelnut exhibits sporophytic self-incompatibility (SSI), whereby compatibility between cultivars is determined by the genotypes of the plants. Incompatibility is determined by a single highly polymorphic locus, with a minimum of two genes, one each for male and female identity. The best-characterized example of SSI is in *Brassica*, which consists of two genes related to pollen-stigma recognition: a female serine/threonine receptor kinase and a cysteine-rich protein that serves as the pollen’s credentials for compatibility interactions (Takasaki et al., 2000; Schopfer et al., 1999). Both proteins co-localize in clusters on the genome containing similar sequences and in the plasma membrane, and are thought to be adapted from pre-existing signaling systems related to pathogen defense (Zhang et al., 2011). To identify SI alleles in hazelnut, the current method is a time-consuming process that requires a library of tester pollens and fluorescence microscopy to visualize pollen germination (Mehlenbacher, 1997); a total of thirty-three SI alleles have been identified thus far with an nine-level dominance hierarchy (Mehlenbacher, 2014). The locus responsible for SI has been mapped to LG 5 (Mehlenbacher et al., 2006). Fine mapping of this locus revealed a region spanning 193 kb and containing 18 predicted genes that differentiate between two SI-alleles, S_1_ and S_3_ (Hill et al., 2021). Previous studies have shown that *Corylus* displays a unique SSI mechanism and is independent of the well-characterized SSI system in Brassica (Hou et al., 2022). Remapping the SI locus will increase the precision of molecular marker development for SI-alleles, enabling further investigation into the genic contributions from parental plants. This will also help reveal the molecular mechanisms involved in *Corylus* SSI and identify candidate genes responsible for SI specificity.

Here we present a chromosome-length haplotype-resolved genome assembly and annotation of ‘Jefferson’. The assembly was produced using Pacific Biosciences HiFi reads and chromosome-scaffolded using high throughput chromosome conformation capture (Hi-C) sequence data. The practical value of this genome assembly is demonstrated by the separation of the two parents into haplotypes at the previously mapped locus for self-incompatibility alleles. Additionally, haplotype separation identified new candidate genes derived from the parent that contributed ‘Gasaway’ EFB resistance, providing insight into the molecular mechanisms of resistance.

## Materials and methods

### Plant material

The *C. avellana* cultivars ‘Jefferson’, and its parents, female ‘OSU 252.146’ and male ‘OSU 414.062’ were used for genome sequencing and assembly. ‘OSU 252.146’ is susceptible to EFB and carries the SI-alleles S_3_ and S_8_, whereas ‘OSU 414.062’ has ‘Gasaway’ resistance and is homozygous for the SI-allele S_1_. Young leaf material was collected from field grown trees in Corvallis, Oregon, USA. Plants were dark-caged for 2-3 days prior to collection, and collected leaves were frozen in liquid nitrogen for Illumina, PacBio, and Hi-C sequencing. For same-day flow cytometry analysis, young leaf tissue was collected in the early morning of May 2020, from a field grown tree of ‘Jefferson’ following leaf budbreak. Flow cytometry reference material was collected the same day from young tomato leaf tissue (*Solanum lycopersicum* L. ‘Stupicke’) from two-week old potted plants grown in the greenhouse.

### DNA extraction, library preparation, and sequencing

PacBio library prep and sequencing were done at the University of Oregon Genomics & Cell Characterization Core Facility (GC3F). High molecular weight genomic DNA was extracted from flash-frozen leaves. Two 8M SMRT cells were sequenced for ‘Jefferson’. To generate HiFi reads, SMRTbell subreads were combined and post-processed with default parameters (CCS.how). Illumina library prep and sequencing of the parents, ‘OSU 252.146’ and ‘OSU 414.062,’ were done at GC3F according to then current Illumina HiSeq 4000 protocols and the iTRU library prep protocol (Glenn et al., 2019) to generate 150 bp paired-end (PE) reads. For Hi-C sequencing, tissue processing, chromatin isolation, and library preparation was performed by Dovetail Genomics (Santa Cruz, CA, USA). The parental libraries were prepared in a manner similar to that of Erez Lieberman-Aiden et al. (2009) and sequenced as 150bp PE reads using the Illumina Hiseq 4000 platform. Illumina reads were demultiplexed using the Stacks v2.0 Beta 10 process_radtags module (Rochette et al., 2019). Demultiplexed reads were checked for quality using FASTQC (version 0.11.5) (Andrews, 2010) and then cleaned by removing adapters, trimming, and quality filtering using the BBTools software suite (Bushnell, 2016); the filterbytile.sh script was used to remove reads associated with low-quality regions of the flow cells containing bubbles, BBDuk was then implemented to trim or remove contaminating iTRU adapters, keep paired reads larger than 130bp, and quality filtering removed reads below Q20.

### Flow cytometry

Flow cytometry was done on ‘Jefferson’ using the propidium iodide (PI) staining technique (Doležel et al., 2005). Solutions of nuclei extraction buffer and staining buffer for PI were prepared using the Cystain^®^ PI kit according to manufacturer protocols (Sysmex, Lincolnshire, IL). Tomato (*Solanum lycopersicum* L. ‘Stupicke’) was used as a reference standard. The 2C DNA content of tomato has been determined to be 1.96 picograms (pg), where 1pg DNA = 0.978 x 10^9^ bp (Doležel et al., 2005). Absolute genomic DNA was calculated by the following formula:

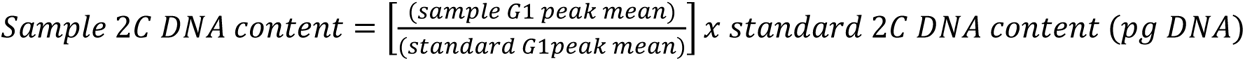

Briefly, sliced leaf squares of tomato and ‘Jefferson’ of equal size (∼0.5cm^2^) were placed in a petri dish together before the addition of 0.5 mL of nuclei extraction buffer. The *C. avellana* samples and tomato standard samples were co-chopped for 30 seconds using a razor blade prior to filtering through a 30 μm nylon-mesh CellTrics^®^ into a 3.5 mL tube. Then, 2 mL of PI staining solution was added to the remaining tissue within the filter. The mixture was incubated at room temperature for 30 minutes inside a Styrofoam cooler to protect against light. Two replicated runs were conducted on different days to account for instrument variation. Stained nuclei were analyzed using a QuantaCyte Quantum P flow cytometer and CyPad software version 1.1. A minimum of 15,000 nuclei counts occurred before the manual gating of G1 sample and standard peaks for each run.

### Genome sequence assembly

An initial Genome size was estimated with a *k-mer* analysis of HiFi reads using Jellyfish (version 2.3.0, RRID: SCR_005491) and the web version of GenomeScope (version 2.0, RRID: SCR_017014) with settings: *k-mer* length of 21 and read length of 15,000 bp (Marçais et al., 2011; Vurture et al., 2017). A haplotype-resolved contig assembly was generated using hifiasm trio-partition algorithm (version 0.16.1-r375, RRID: SCR_021069) (Cheng et al., 2021). First, individual *k-mer* counts of parental Illumina reads of the parents ‘OSU 252.146’ and ‘OSU 414.062’ were acquired using Yak (version 1.1) as input evidence for hifiasm trio binning. The Arima Hi-C mapping pipeline was followed to generate mapped Hi-C reads (Github.com/ArimaGenomics/mapping_pipeline). YaHs (version 1.1, RRID: SCR_022965) was run independently on both haplotype assemblies produced by hifiasm with their respective Hi-C aligned, read-name sorted bam file (Zhou et al., 2022). A Hi-C contact map was generated for each respective haplotype. Contigs were combined and gapfilled using Juicebox (version 1.11.08, RRID: SCR_021172) (Durand et al., 2017); finalized Hi-C contact maps were curated by Hudson Alpha (Huntsville, AL, USA), using an unpublished Hi-C scaffolding and alignment tool that oriented ‘Jefferson’ chromosomes based on the ‘Tombul’ genome pseudo-chromosomal scaffolds (Lucas et al., 2020). To verify haplotype assignment accuracy, parental reads were realigned to each haplotype assembly. Final assembly metrics were generated by QUAST (version 5.0.0, RRID: SCR_001228) (Mikheenko et al., 2018). Assembly completeness was assessed with BUSCO (version 5.4.6, RRID: SCR_015008) in genome mode, using the Embryophyta odb10 dataset (Manni et al., 2021). The quality of assembling repetitive genomic regions were assessed using the long terminal repeat (LTR) assembly index (LAI); this pipeline was composed of LTRharvest within GenomeTools (version 1.6.1, RRID: SCR_016120), LTR_FINDER (version 1.2, RRID: SCR_015247), and LTR_retriever (version 2.9.4, RRID: SCR_017623) using suggested default parameters to predict and combine likely full length candidate LTR-RTs (retrotransposons) (Ou et al., 2018). Calculation of the LAI index was based on the formula: LAI= (intact LTRs/total LTR length) x 100.

### Structural gene annotation

Gene prediction and annotation was facilitated by Illumina transcriptome data from the following sources: 1) ‘Jefferson’ style, bark and leaf tissue, *C. avellana* ‘Barcelona’ catkins, whole seedling of ‘OSU 954.076’ x ‘OSU 976.091’ including root tissue (Rowley et al., 2012; Sathuvalli, unpublished); and 2) leaf bud tissue from *C. avellana* ‘Tombul’, ‘Çakildak’, and ‘Palaz’, publicly available from the National Center for Biotechnology Information (SRA: PRJNA316492) (Kavas et al., 2020). The resulting set of reads putatively representing *C. avellana* was ∼423 million PE 150bp RNA-seq reads. Similarly, a protein set consisting of 61,590 annotated proteins was curated from a previous unpublished version 3 ‘Jefferson’ genome assembly and *C. avellana* ‘Tombul’ (Lucas et al., 2021). Gene annotation was performed for both ‘Jefferson’ haplotype assemblies. To create a repeat library of transposable element families, a RepeatModeler (RRID: SCR_015027) families set was concatenated with the haplotype-resolved chromosome-level assemblies of ‘Jefferson’ and six other OSU *C. avellana* accessions that were trio-assembled using the same methods as ‘Jefferson’ but without chromosome scaffolding (unpublished). Low complexity DNA sequences and repetitive regions were soft masked prior to gene annotation using the default parameters of RepeatMasker (version 4.1.0, RRID: SCR_012954). Structural annotations of protein-coding genes were identified using the gene prediction software AUGUSTUS, GeneMark-ES/EP+, and GenomeThreader, integrated by BRAKER1 and BRAKER2 (RRID: SCR_018964) (Stanke et al., 2006a,b, 2008; Li et al., 2009; Barnett et al., 2011; Gremme, 2013; Lomsadze et al., 2014; Buchfink et al., 2015; Hoff et al., 2016, 2019; Brůna et al., 2020, 2021). First, BRAKER1 used a unique .bam file generated from the splice-aware aligner Hisat2 (Kim et al., 2019), of the previously described RNA-seq set aligned to each haplotype assembly. Second, BRAKER2 was run using the AUGUSTUS *Arabidopsis thaliana* training set and gene structures were predicted via spliced alignments with AUGUSTUS ab-initio and GenomeThreader integration on each masked haplotype genome using the combined protein dataset previously described. Gene predictions of the respective BRAKER1 and BRAKER2 haplotype runs were assessed for quality, deduplicated, and combined using TSEBRA with default settings (Gabriel et al., 2021).

To further improve this original gene annotation set, BRAKER3 was used (Gabriel et al., 2023). A new masked genome was generated for both haplotype assemblies using EDTA (version 2.1.0, RRID: SCR_022063) (Ou et al., 2019) with parameters: --anno 1 --cds --sensitive including the respective coding sequences and gene locations generated by the BRAKER1/BRAKER2 pipeline. Finalized gene prediction sets were produced using BRAKER3 that included soft-masked genomes, a curated Viridiplantae ODB11 protein set consisting of roughly 5.3 million proteins, and the previously described RNA-seq dataset.

BRAKER3 outputs were used as input for TSEBRA, with the -k parameter, to enforce and recover potential missing genes and transcripts produced by the BRAKER1/BRAKER2 pipeline.

### Functional gene annotation

Both haplotype annotation sets from TSEBRA were subject to predictive functional analysis using the transcript set within OmicsBox (version 3.0); the OmicsBox pipeline included CloudBLAST using BLASTx, InterPro, GO Merge, GO Mapping, and GO Annotation plus validation (Altschul et al., 1990; Götz et al., 2008; Paysan-Lafosse et al., 2022). Completeness of the predicted annotation sets was assessed using BUSCO --protein mode, inputting translated amino-acid sequences derived from CDS of gene transcripts and the Embryophyta odb10 dataset. To assess long-range structural variation between haplotype assemblies, translocations, inversions, and copy number variation were identified using minimap2 (version 2.23-r1111, RRID: SCR_018550) (Li H., 2018), and SyRI (version 1.6.3, RRID: SCR_023008) and visualized by plotsr (Goel et al., 2019, 2022). Conservation of putative high confidence homologs between assemblies were compared using Orthofinder (version 2.5.4, RRID: SCR_017118) (Emms and Kelly, 2019).

### Identification of candidate genes for EFB resistance and self-incompatibility

To identify potential disease resistance gene homologs, the amino acid sequence of annotated protein-coding genes from each assembly were queried against the Plant Resistance Gene Database (version 3.0) using DRAGO2-api (Osuna-Cruz et al., 2018). DNA alignments of previously identified RAPD and SSR marker sequence fragments, BAC-end libraries, and annotated protein-coding genes from ‘Jefferson’ were aligned to the new genome assemblies using minimap2 (Heng Li, 2018). Marker locations were secondarily assessed for off-target allele-size amplification and multimapping by *in silico* PCR using each marker’s corresponding primer pair mapped against the Jefferson V4 haplotype 1 and 2 genomes, allowing for 1-2 mismatches per primer pair. A multiple sequence alignment of the translated candidate R-genes from each haplotype was generated with MUSCLE (version 5.1.0, RRID: SCR_011812) using default settings (Edgar, 2021). A phylogenetic tree of these sequences was created using the neighbor joining tree (BLOSUM62) calculation in JalView (Waterhouse et al., 2009). MEME software (version 5.4.1, RRID: SCR_001783) was utilized to identify conserved subdomains among the putative R-gene candidate proteins using the settings: -mod anr -nmotifs 10 -protein (Bailey et al., 2009).

In a similar approach, genes involved in self-incompatibility were remapped to both haplotype assemblies using previously identified fine-mapped markers and gene sets (Hill et al., 2021). These markers and genes served as query evidence in BLASTn/BLASTp searches of both haplotype assemblies. A multiple sequence alignment of the identified proteins of interest in each haplotype was generated using MUSCLE and visualized using the neighbor joining tree (BLOSUM62) within JalView. The complete genome assembly and annotation pipeline are summarized (Supplemental Figure S1).

## Results and discussion

### Genome assembly

A combined total of 3.6 million PacBio HiFi reads with an average length of 15,597 bp were generated from two 8M SMRT cells, resulting in 56.8 Gb of sequence data (∼147x genome coverage) (Supplementary table S1). For the two parents, ‘OSU252.146’ and ‘OSU414.062’, 295 and 218 million PE 150 bp Illumina reads were generated, yielding 44 Gb (115x coverage) and 32 Gb (85x coverage), respectively (Supplemental table S1). These reads were used to generate hifiasm trio binned haploid genome assemblies spanning 385,825,918 bp and 372,534,284 bp, containing 663 and 229 contigs for haplotype 1 and 2, with N50s of 23.4 Mb and 22.5 Mb, respectively (Table 1).

The hifiasm haplotype assemblies were used as inputs to the chromosome scaffolding process. Hi-C sequencing of ‘Jefferson’ generated ∼428 million PE 150 bp reads, for a total yield of ∼64.6 Gb (168x coverage, Supplemental table S1). The resulting ‘Jefferson V4’ Hi-C scaffolded genome assemblies of each haplotype consisted of 11 pseudo-chromosomal scaffolds. The chromosome-level assemblies spanned a total length of 349,702,244 bp and 352,009,510 bp for haplotype 1 and haplotype 2, an N50 of 32.5 Mb and 32.4 Mb (Table 1, Supplemental table S2). The Hi-C interaction matrix clearly differentiated between individual chromosomes in both haplotypes (Figure 1A, 1B). Alignment of parental reads to each genome assembly haplotype showed that the majority of reads from ‘OSU 252.146’ aligned to haplotype 2, whereas the majority of reads from ‘OSU 414.062’ aligned to haplotype 1 (Supplemental table S3). BUSCO results in genome mode showed that both chromosome-level haplotype genome assemblies were of high, comparable quality and captured >97% of conserved genes in the Embryophyta dataset (Table 2).

**Figure 1A.**
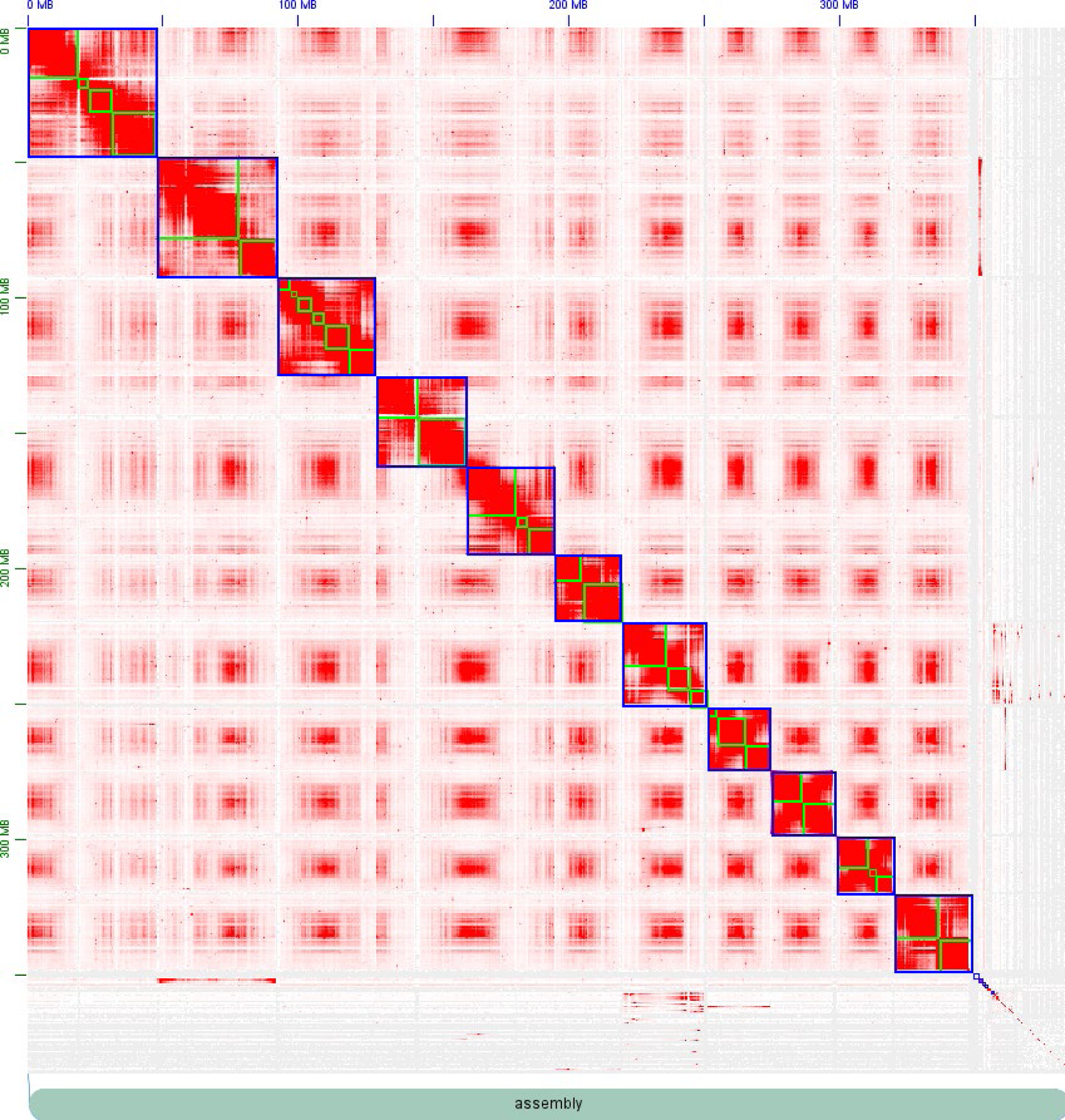
Hi-C interaction matrix for the ‘Jefferson’ (*C.* avellana) haplotype 1 assembly. On the X and Y-axes is the distance in the genome assembly (Mb), the green squares represent contigs that are scaffolded within the blue square, which represent a chromosome. The red indicates chromatin interaction loci which are most abundant within chromosomes. The grey space in the lower right represent unaligned contigs which did not have sufficient Hi-C mapping depth to be incorporated into chromosomal scaffolds.

**Figure 1B.**
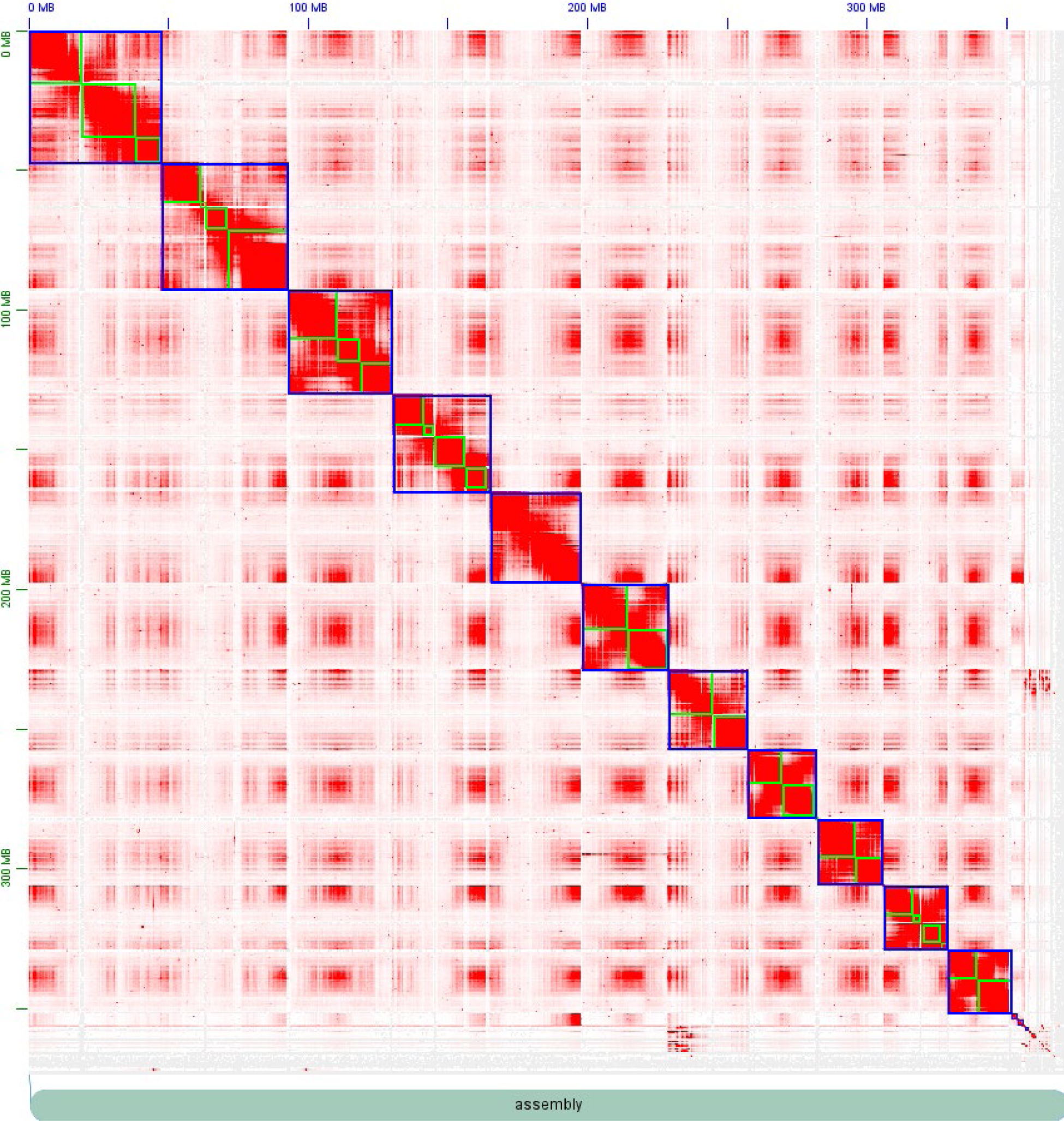
Hi-C interaction matrix for ‘Jefferson’ (*C. avellana*) haplotype 2 assembly.

**Table 2.**
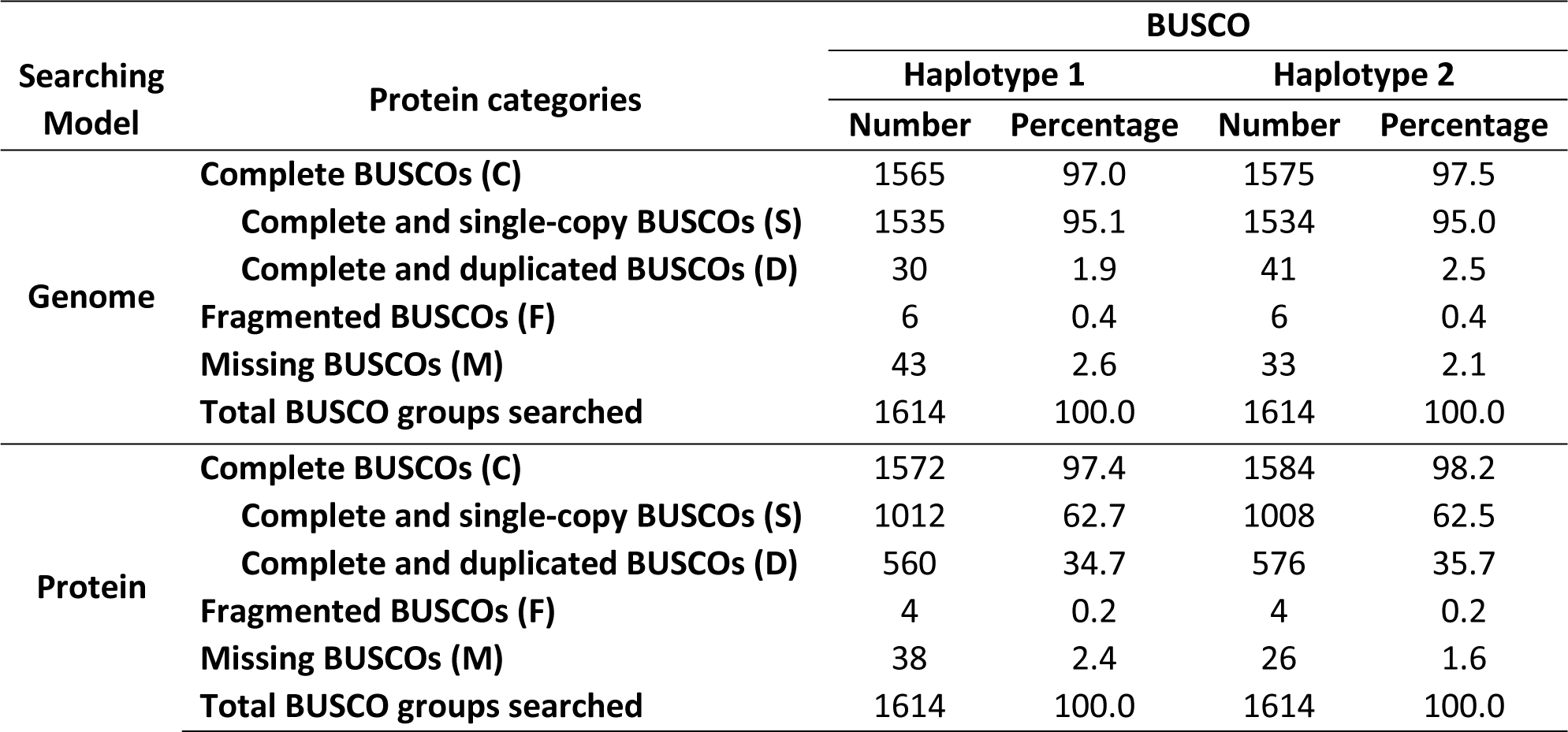
Assessment of genome completeness in ‘Jefferson’ haplotypes using BUSCO.

### Genome size estimation

Flow cytometry was used to estimate a 1C genome size of ‘Jefferson’ of 365.65 Mb (1C = 0.37 pg). This estimate is slightly smaller than a previously reported estimate of ‘Jefferson’ (370 Mb) (Rowley et al., 2018) and the reported range of other cultivars and diploid species in the subgenus *Corylus* (C*. cornuta*, *C. colurna*), which was between 1C = 0.41 - 0.43 pg (Bai et al., 2012; Vallès et al., 2014). PacBio HiFi reads of ‘Jefferson’ were also input to GenomeScope to provide a secondary genome size estimate and heterozygosity of 274.8 Mb and 1.54%, respectively (Supplementary Figure S2). The *k-mer* based estimate is significantly less than the flow cytometry estimate, likely due to limitations of the algorithm in accounting for long-read length and high heterozygosity. The chromosome-resolved assemblies were 4% smaller than the flow cytometry prediction.

### Linkage map of ‘Jefferson’

The first available *Corylus avellana* linkage map was constructed using random amplified polymorphic DNA and simple sequence repeat (SSRs)markers segregating in an F1 mapping population derived from a cross between ‘OSU 252.146’ and ‘OSU 414.062’, the same population from which ‘Jefferson’ was selected (Mehlenbacher et al., 2004). Since then, this linkage map has been improved by additional SSRs and data from a bacterial artificial chromosome (BAC) library (Sathuvalli et al., 2017; Mehlenbacher and Bhattarai, 2018). To assign the linkage groups to pseudo-chromosomal scaffolds, 18 RAPD, 874 microsatellite, 4,100 paired BAC-ends with proper insert size, and 15,000 biallelic SNP marker sequence fragments were aligned to both Jefferson haplotypes using minimap2 (Li H., 2018), and compared to previous linkage mapping designations (Koma Komaei, 2020). Both haplotypes were successfully assigned the same linkage group for each corresponding pseudo-chromosomal scaffold and renamed appropriately.

### Synteny of ‘Jefferson’ haplotypes

The ‘Jefferson’ haplotype assemblies showed a high degree of synteny (Figure 2). Differences in length between pseudo-chromosome haplotypes ranged from ∼16,000 bp (chromosome 6) to ∼2.5 Mb (chromosome 5); most scaffolds representing homologous chromosomes differed in length by an average of ∼892 kb. Between haplotypes there were three large scale translocations (chromosome 2, 7, and 9), two inversions (chromosome 5 and 6), and several small duplications, translocations, and gaps. The most notable of non syntenous regions were three large scale translocations on chromosomes two, seven, and nine, comprising total lengths of 14 Mb, 13 Mb, and 9.7 Mb, respectively (Supplemental table S4). Despite nearly 93% of the haplotype assemblies mapping to one another, 33% of the alignments were categorized as having high divergence (Supplemental table S5). Past cytological work has categorized three chromosome sizes, with two homologous pairs being large, five medium, and three small (Falistocco and Marconi, 2013). Translocations have also been observed in *Corylus* (Salesses and Bonnet, 1988). Reciprocal translocations are thought to frequently confound genetic map generation for many hazelnut populations (Lunde et al., 2006; Bhattarai et al., 2017; Marioni et al., 2018), and are hypothesized to be the result of cytogenetic abnormalities, such as irregular chromosomal migration during cell division, or nondisjunction during microsporogenesis or megasporogenesis (Lagerstedt, 1977). Mono-, bi-, and multi-valent chromosome pairings have been observed frequently in *Corylus* spp. and their hybrids (Woodworth, 1929; Kasapligil, 1968); this suggests that unequal crossover events may be common, especially when diverse germplasm is used. However, it is also possible these apparent translocations are errors from orienting the ‘Jefferson’ Hi-C alignment against ‘Tombul’.

**Figure 2.**
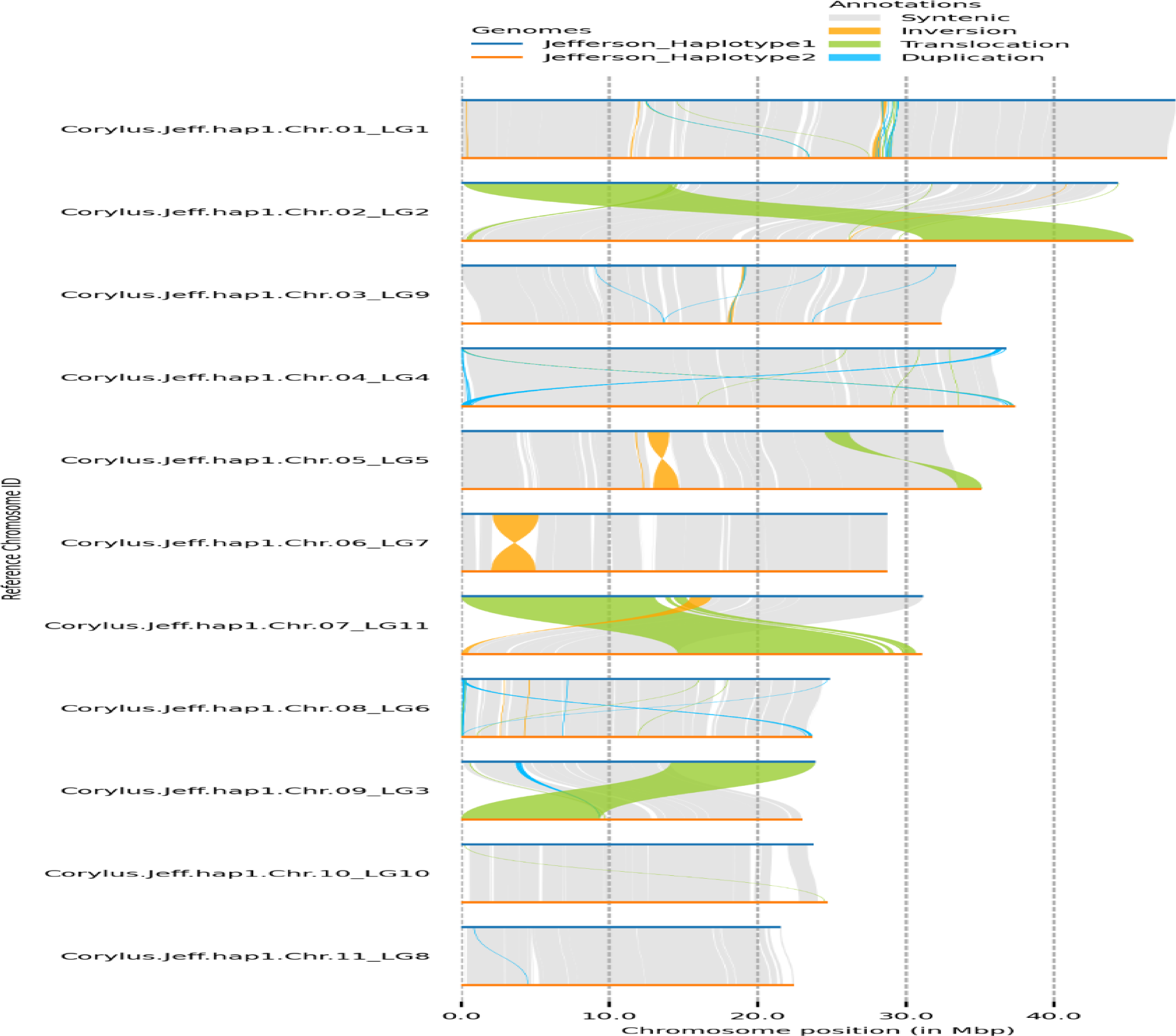
Synteny plot of the two ‘Jefferson’ chromosome-resolved haplotype assemblies. Pseudo-chromosomal scaffolds of each haplotype were aligned to each other, and labelled on the Y-axis with the chromosome ID and related linkage group. The X-axis shows the chromosome size in Mbp. Chromosomes of haplotypes 1 and 2 are displayed as blue and orange lines, respectively. Grey shading represents complete synteny between genomic positions, yellow represents an inversion, green represents a translocation, and light blue represents a duplication.

### Characterization of repeats

Prior to annotating protein-coding genes, genome repeat identification and masking was performed on the chromosome-level haplotype assemblies. The proportion of repeats and unknown elements identified in the initial RepeatModeler and RepeatMasker runs for the ‘Jefferson’ haplotypes was higher than those reported for other *C. avellana* cultivars and *Corylus* species, with ∼65% of bases being masked. The high proportion of LTRs identified suggested potentially erroneous repeat calls that were introduced by the large concatenated LTR families dataset. By rerunning the analysis using EDTA, a more stable view of LTRs was obtained, with 38.26% and 35.29% of repeats masked for haplotype 1 and 2 (Supplemental table S6, S7). Class I retroelements made up 46-54% of all repeats identified for haplotype 1 and 2, respectively. *Gypsy* superfamilies were nearly double those of *Copia*, which is opposite of what has been previously reported in *C. avellana ‘Tombul’* but on par with *C. avellana ‘Tonda Gentile delle Langhe’* and Silver birch (*Betula pendula*) ‘SB1’ (Lucas et al., 2021; Pavese et al., 2021; Salojärvi et al., 2017). Nearly 20% of the total repeat length identified in either haplotype had LTRs categorized as ‘unknown.’ The most significant difference observed between repeat elements of the haplotype assemblies was a doubling of the loosely-defined annotated “repeat_region”, with 21 Mb and 9.5 Mb for haplotype 1 and 2, respectively. LAI analysis of haplotype 1 (LAI=16.9) and haplotype 2 (LAI=16.2), indicates that the repetitive and intergenic sequence space is of reference genome quality and a significant improvement from *‘Tombul’* (LAI=8.76) (Supplementary Figure S3).

### Structural and functional gene annotation

A total of 32,431 and 33,159 protein-coding genes were identified in haplotypes 1 and 2, respectively, and when considering alternative isoforms, these numbers increased to 48,832 and 50,663 coding transcripts, respectively. The protein-coding genes of both haplotype assemblies had an average length of 3,653/3,695 bp, with an average of 3.5 introns per longest isoform and median intron and exon lengths of 232 and 138 bp, respectively. For haplotypes 1 and 2, 21,201/21,354 (∼64%) of genes had no alternative isoforms, 7,767/8,089 (∼24%) had one alternative isoform and 3,453/3,716 (∼11%) had two or more isoforms. For each haplotype’s predicted gene set, >97% of *C. avellana* genes were complete BUSCOs for the ODB10 Embryophyta gene families (Table 2). Approximately 35% of highly conserved BUSCO genes were predicted as complete-duplicated, likely due to alternative transcripts.

For haplotype 1, functional annotation analyses assigned GO terms and InterPro domains to 24,369 (72.7%) of transcripts. For the remaining transcripts in haplotype 1, 3,907 (11.4%) had no blast hits, 3,605 (10.8%) had only blast hits, 1,666 (5%) were identified with GO mapping. Similarly for haplotype 2, 24,932 (72.5%) of transcripts were assigned GO terms and InterPro domains. Of the remaining transcripts in haplotype 2, 3,907 (11.4%) had no blast hits, 3,725 (10.8%) had only blast hits, and 1,815 (5.3%) of transcripts were GO mapped (Supplemental Figure S4, S5). OrthoFinder was used to further characterize and assess conservation between predicted gene sets of each haplotype assembly. Of the combined 99,495 transcripts from haplotype 1 and 2, 96,193 (96.7%) were placed in a total of 31,779 orthogroups, with only 4,618 (4.6%) of genes being categorized as unique to a haplotype. To assess the overall distribution of disease resistance genes, DRAGO2 identified 3,620 and 3,659 putative genes with resistance-like domains for haplotype 1 and haplotype 2 assemblies. The majority of these genes identified by DRAGO2 were receptor-like kinases and proteins (∼25%), with a small fraction being identified as NBS-LRRs (∼10%) (Supplement table S8).

### Potential candidate genes for self-incompatibility

The locus for pollen-stigma incompatibility was fine-mapped by Hill et al. 2021, who identified 18 genes within a 193.5 kb region on linkage group 5 that were associated with SI alleles S_1_ and S_3_. To remap the SI locus, BLASTn was used to align genes from the previous assembly to both chromosome-resolved haplotype assemblies of ‘Jefferson.’ BLASTn searches returned twelve genes with 100% identity to the S_1_ allele among the newly predicted genes in haplotype 1, chromosome 5. In chromosome 5 of haplotype 2, eleven genes with 100% identity to the S_3_ allele were identified. Multiple genes that were previously identified as candidates for SI interactions in *Corylus*, PIX7 (Putative interactor of XopAC_7_) and MIK2 (*MDIS_1_-interacting receptor like kinase*) were also found in both Jefferson haplotypes. Haplotype 1 contained two copies of PIX7 and eight copies of MIK2, whereas haplotype 2 contained three copies of PIX7 and five copies of MIK2. The SI-locus occupied 86.6 kb in haplotype 1 and 222 kb in haplotype 2. The phasing of alleles within the chromosome 5 SI locus agrees with the previous fine mapping results showing that ‘OSU 252.146’ contributes S_3_ to ‘Jefferson’, and is represented in the haplotype 2 assembly, whereas ‘OSU 414.062’ which contributed S_1_ to ‘Jefferson’, is represented in the haplotype 1 assembly.

The similarity of PIX7 and MIK2 candidates was assessed using OrthoFinder, which assigned these genes to seven orthogroups. All seven PIX7 homologs were assigned to three orthogroups, whereas the majority of MIK2 homologs were assigned to a single orthogroup. This suggests that putative PIX7 and MIK2 candidate gene copies are highly conserved, but there may be some variation in protein subdomains that lead to the identification of multiple orthogroups. Indeed, of the eighteen genes identified as PIX7 or MDIS-1 homologs, all were variable in total length (Table 3). Recent studies have shown that in *Brassica*, the most well characterized SSI system, a small RNA is crucial for inducing methylation of recessive SI allele, in order to induce compatibility (Yasuda et al., 2021). When considering the large number of SI-alleles in *Corylus* (33 to date), it is possible that unannotated sRNA(s) are acting upon different variants of PIX7 or MIK2 to establish allelic dominance. Additional genomes of other *Corylus* cultivars with confirmed SI-alleles will be needed to verify differences in SI-alleles and putative candidate genes to further elucidate the complex molecular mechanism driving SSI and allelic hierarchy in *Corylus*.

**Table 3.** *Corylus avellana* ‘Jefferson’ self-incompatibility homologs identified in the self-incompatibility region of both haplotypes of chromosome 5 (LG 5).

| *Corylus avellana* gene | Amino acid length (bp) | Function |
| --- | --- | --- |
| Hap1_g18435 | 513 | probable serine/threonine protein kinase PIX7 |
| Hap1_g18437 | 695 | MDIS1 interacting receptor like kinase 2 like |
| Hap1_g18438 | 328 | MDIS1 interacting receptor like kinase 2 like |
| Hap1_g18439 | 937 | MDIS1 interacting receptor like kinase 2 like |
| Hap1_g18441 | 357 | MDIS1 interacting receptor like kinase 2 like |
| Hap1_g18442 | 767 | MDIS1 interacting receptor like kinase 2 like isoform |
| Hap1_g18443 | 1,056 | MDIS1 interacting receptor like kinase 2 like isoform |
| Hap1_g18444 | 112 | probable serine/threonine protein kinase PIX7 |
| Hap1_g18445 | 177 | MDIS1 interacting receptor like kinase 2 like isoform |
| Hap1_g18450 | 787 | MDIS1 interacting receptor like kinase 2 like isoform |
| Hap2_g19113 | 477 | probable serine/threonine protein kinase PIX7 |
| Hap2_g19115 | 417 | MDIS1 interacting receptor like kinase 2-like |
| Hap2_g19117 | 937 | MDIS1 interacting receptor like kinase 2-like |
| Hap2_g19118 | 182 | probable serine/threonine protein kinase PIX7 |
| Hap2_g19119 | 950 | MDIS1 interacting receptor like kinase 2-like |
| Hap2_g19124 | 793 | MDIS1 interacting receptor like kinase 2-like |
| Hap2_g19138 | 1,296 | MDIS1-interacting receptor like kinase 2-like |
| Hap2_g19148 | 513 | probable serine/threonine protein kinase PIX7 |

### Potential candidate genes for EFB resistance in hazelnut

In ‘Jefferson,’ EFB resistance is derived from ‘Gasaway’ and is conferred by a dominant allele at a single locus that has been mapped between RAPD markers 152-800 and 268-580 on linkage group 6 (Mehlenbacher et al., 2006). Recent QTL (Quantitative Trail Loci) mapping in *C. americana* x *C. avellana* mapping populations associated LG6 EFB resistance in *C. avellana* cv. ‘Tonda di Giffoni’, with SNP 93212 (Lombardoni et al., 2022). Aligning the associated paired-end sequences from SNP 93212 to ‘Jefferson’ V4 haplotype 1 placed the QTL peak 20 kb upstream from the markers most closely associated with EFB resistance, and within BAC contig 43F13 in the fine-mapped region defined by Sathuvalli et al. (2017). When mapping the Sanger sequence of CC875206.1 W07-365 (365 bp), the RAPD marker originally extracted from the PCR band associated with W07 ‘Gasaway’ resistance, the sequence is repeated 3 times in this region in both haplotypes of ‘Jefferson;’ however, the sequence is truncated by ∼60 bp in haplotype 2 and spans an additional 100 kb in chromosomal space. Mapping the original Illumina reads from BAC 43F13 to both haplotypes revealed haplotype 1 as the source of the BAC contig and clearly defined the region coinciding with the associated BAC-end markers. The higher percentage of Illumina reads aligning to haplotype 1 from EFB-resistant parent ‘OSU 414.062’, provides additional support for an EFB-resistance model with R-gene contributions derived from ‘Gasaway’ present in haplotype 1 only.

Functional annotation of the ‘Jefferson’ EFB resistance region on haplotypes of chromosome 8 (LG 6) identified several probable receptor-like kinases and putative disease resistance genes. On haplotype 1, a region of approximately 125 kb contained five CNLs identified by DRAGO2 but eight genes with functional descriptions relating to “RGA” (Resistance Gene Analog). On haplotype 1, Hap1_g26572 and Hap1_g26573 were identified as having homology to RGA3 and a short 232aa RGA2-like isoform, respectively. Six other putative resistance genes were identified in haplotype 1, including a long 1,116 aa copy of disease resistance RGA2-like isoform in Hap1_g26576, three copies of RGA3 in Hap1_g26579, Hap1_g26581, and Hap1_g26582, and two copies of RGA4 in Hap1_g26580 and Hap1_g26583. Similarly, haplotype 2 contained fourteen genes with functional descriptions related to “RGA3” and “RGA2-like isoform” (Table 4), but only eleven were identified as CNLs by DRAGO2. None of the R-genes from haplotype 1 had a 100% match to haplotype 2 R-genes. In Figure 3, the genomic location and orientation of the putative EFB R-gene candidates on chromosome 8 (LG6) are depicted for both haplotypes, showing that RGA3 homologs are closely linked to an RGA2-like isoform and an RGA4 homolog on haplotype 1, whereas R-gene candidates on haplotype 2 are identified as only RGA3 and one as RGA2-like isoforms, all ranging in distance from one another by 20-60 kb.

**Figure 3.**
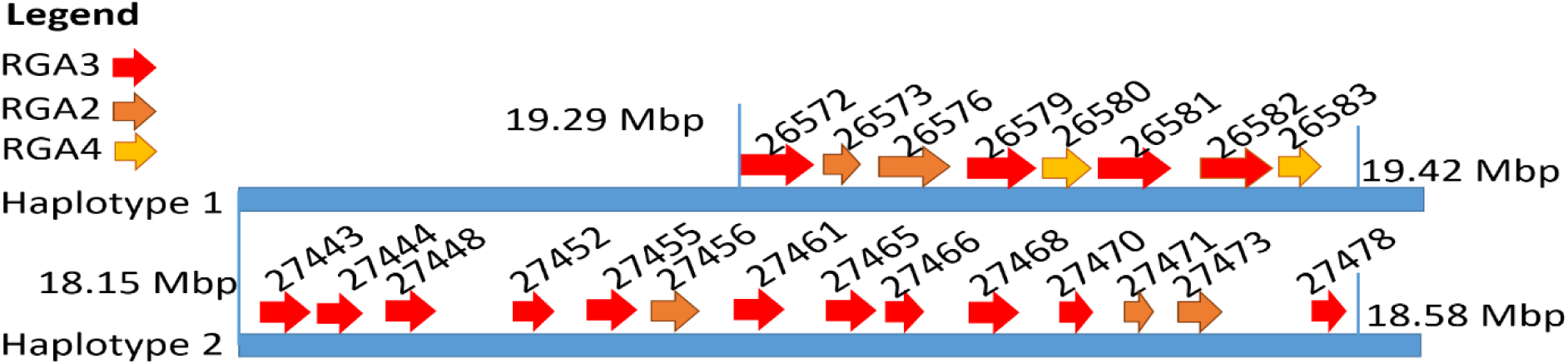
Putative EFB R-gene candidates (RGA-homologs) plotted on chromosome 8 of both haplotypes. Red arrows represent RGA3 homologs, orange arrows represent RGA2 isoform-X2 homologs and yellow arrows represent RGA4 homologs. The gene ID for each respective homolog is listed above the arrow where haplotype 1 represents Hap1_g and haplotype 2 represents Hap2_g. Denoted as vertical lines in Mb are the start and stop positions of the R-gene cluster.

**Table 4.** *Corylus avellana* ‘Jefferson’ candidate EFB R-gene homologs identified in the ‘Gasaway’ resistance region locus on chromosome 8 (linkage group 6) of both haplotypes.

| *Corylus avellana* gene | Amino acid length (bp) | Function |
| --- | --- | --- |
| Hap1_g26572 | 1,213 | putative disease resistance protein RGA3 |
| Hap1_g 26573 | 237 | disease resistance protein RGA2-like isoform X2 |
| Hap1_g 26574 | 224 | Cysteine-rich repeat secretory protein 38 |
| Hap1_g 26576 | 1,116 | disease resistance protein RGA2-like isoform X2 |
| Hap1_g 26579 | 891 | putative disease resistance protein RGA3 |
| Hap1_g 26580 | 323 | putative disease resistance protein RGA4 |
| Hap1_g 26581 | 1,191 | putative disease resistance protein RGA3 |
| Hap1_g 26582 | 849 | putative disease resistance protein RGA3 |
| Hap1_g 26583 | 274 | putative disease resistance protein RGA4 |
| Hap1_g 26585 | 230 | Cysteine-rich repeat secretory protein 38-like |
| Hap2_g27443 | 1,215 | putative disease resistance protein RGA3 |
| Hap2_g27444 | 1,164 | putative disease resistance protein RGA3 |
| Hap2_g27448 | 1,145 | putative disease resistance protein RGA3 |
| Hap2_g27452 | 1,159 | putative disease resistance protein RGA3 |
| Hap2_g27455 | 1,159 | putative disease resistance protein RGA3 |
| Hap2_g27456 | 1,150 | disease resistance protein RGA2-like isoform X2 |
| Hap2_g27461 | 1,178 | putative disease resistance protein RGA3 |
| Hap2_g27465 | 1,145 | putative disease resistance protein RGA3 |
| Hap2_g27466 | 847 | putative disease resistance protein RGA3 |
| Hap2_g27468 | 1,159 | putative disease resistance protein RGA3 |
| Hap2_g27470 | 472 | putative disease resistance protein RGA3 |
| Hap2_g27471 | 725 | disease resistance protein RGA2-like isoform X2 |
| Hap2_g27473 | 1,150 | disease resistance protein RGA2-like isoform X2 |
| Hap2_g27478 | 473 | putative disease resistance protein RGA3 |

RGA4 has been characterized as an auto-inducer of immune response to the fungal disease rice blast caused by *Magnaporthe oryzae*, whereby RGA4 is tightly linked with RGA5, with the encoded proteins interacting as a homo and hetero dimer, such that both are required for resistance (Césari et al., 2014). Research suggests that the presence of an integrated heavy metal associated (HMA) domain within RGA5 mimics the pathogen effector target as a “decoy”, and upon direct binding to the effector, a signal is transduced to RGA4, relieving RGA4 repression and initiating an immune response (Xi et al., 2022). Heavy metal-associated isoprenylated plant proteins (HIPPs) in rice (*Oryza sativa*) contain HMA domains, and have been identified as putative effector hubs (Bentham et al., 2020; Maidment et al., 2021) as HIPPs have been shown to be the target of multiple fungal effector proteins, having a greater binding affinity to *M. oryzae* AVR-Pik variants than the integrated HMA domains present in rice CC-NLR resistance genes *Pik-1* and *Pik-2* (Maidment et al., 2021). Importantly, HMA domain variants have been shown to perceive new effectors (Césari et al., 2022). On haplotype 1, the genes Hap1_g26587 and Hap1_g26589 were given the functional description “heavy metal-associated isoprenylated plant protein 47” and are located 19 kb and 43 kb upstream, respectively, of the closest RGA4 on the minus strand. Conversely, on haplotype 2, four HMA genes with the same descriptor (Hap2_g27459, Hap2_g27477, Hap2_g27482, and Hap2_g27484) were identified. These genes ranged from 20-72 kb away from the nearest putative RGA3 gene. HIPP genes of haplotypes 1 and 2, respectively, share high identity with minimal amino acid substitutions among each other. Performing a BLASTp of these predicted proteins against the entire protein set of both haplotypes resulted in matches with other predicted HIPPs, with no homology to suggest that the nearby RGA cluster has a unique synonymous integrated HMA domain like that in rice.

In recent years it has become apparent that cysteine-rich receptor-like secreted proteins (CRRSPs) have crucial involvement in plant-fungal pathogen interactions (Zeiner et al., 2023). *Gnk2* from ginkgo (*Ginkgo biloba*) and two maize (*Zea mays*) proteins, *AFP1* and *AFP2*, bind to mannose during the defense response against fungal pathogens (Miyakawa et al., 2014; Ma et al., 2018). Mannose and its reduced sugar alcohol, mannitol, are independently important to both host plant and fungal pathogen metabolism and signaling during plant growth and pathogen invasion (Patel and Williamson, 2016). CRRSPs have also been shown to be directly involved in fungal pathogen recognition as co-receptors for pathogen effectors (Wang et al., 2023). Recently *TaCRK3,* a CRRSP in wheat, was revealed to inhibit mycelial growth *in vitro* (Guo et al., 2021). Five genes were given the functional description “cysteine-rich repeat secretory protein 38”: two in haplotype 1, Hap1_g26574 and Hap1_g26585, and three in haplotype 2, Hap2_g27475, Hap2_g27480, and Hap2_g27457. To further investigate similarity between these CRRSPs, we performed a BLASTp and used MUSCLE to generate a neighbor-joining tree in JalView (Figure 4). The haplotype 1 gene Hap1_g26574 has two transcripts, with .t1 containing a 20 bp deletion at the 5’ end; the two transcripts have an 86% and 88% similarity to the haplotype 1 gene (Hap1_g26585) and the haplotype 2 genes, respectively, whereas all haplotype 2 genes are 100% identical. These genes contained an extracellular domain composed of two DUF26 (domain of unknown function 26) motifs, but notably lacked an intracellular serine/threonine kinase domain and transmembrane domain (Figure 5).

**Figure 4.**
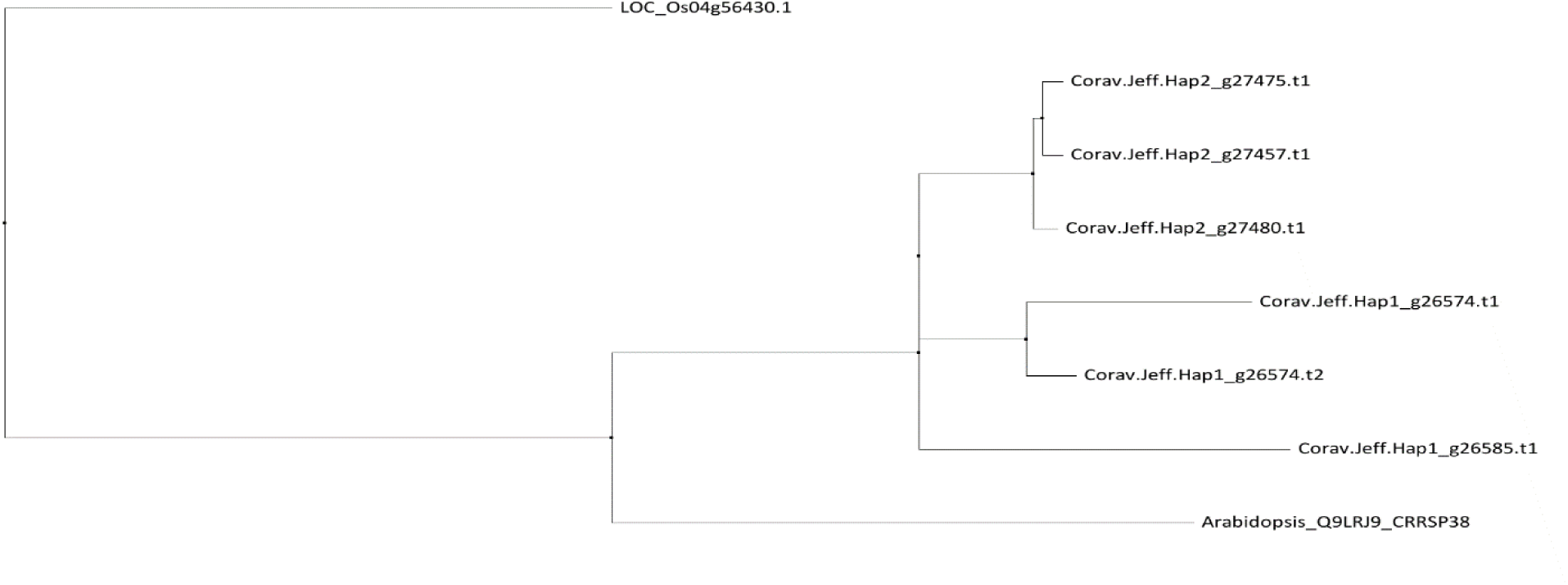
Neighbor joining tree of seven cysteine-rich secretory proteins (CRSPs) within the EFB R-gene region of both haplotypes with *Arabidopsis* and rice (*RCR3*) homologs aligned by MUSCLE.

**Figure 5.**
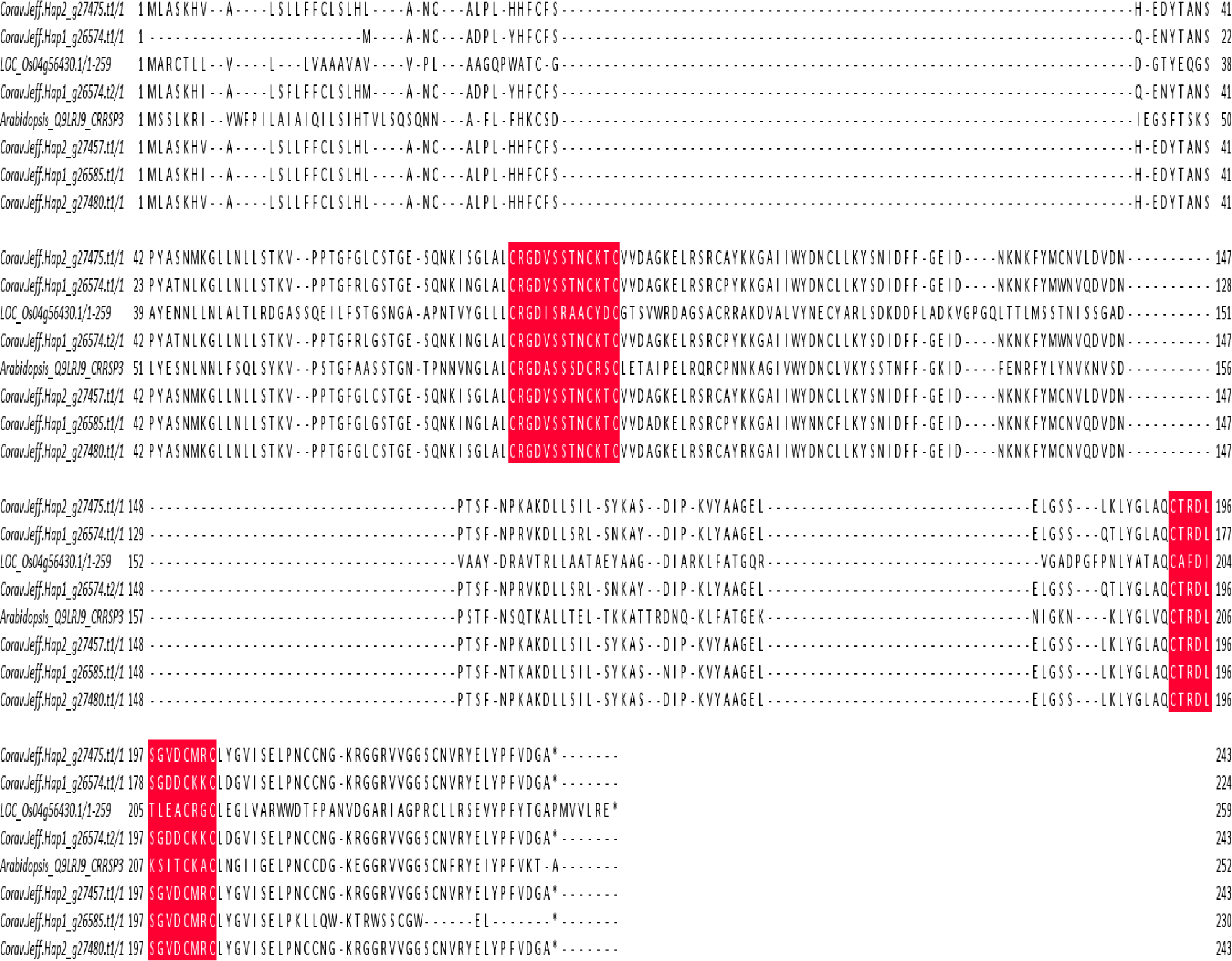
Amino-acid sequence alignment of all Cysteine-rich repeat secretory protein-38 in *C. avellana ‘Jefferson’ and* a homolog from Arabidopsis and Rice (*RMC*). The numbers on the right side indicate the positions of the residues in the corresponding protein. Red shading indicates the conserved motif of the DUF26 domain C-X8-C-X2-C.

Despite identifying candidate EFB resistance genes on haplotype 1, the overall similarity between these genes and haplotype 2 R-genes makes it challenging to determine whether one or several resistance genes are involved in the activation of ‘Gasaway’ resistance. It remains to be determined how the unique CRRSP (Hap1_g24474) is involved in processes of pathogen detection and downstream signaling response with close proximity to numerous NBS-LRRs. Thus, it appears that the uncharacterized disease resistance signaling pathway of ‘Gasaway’ involves NBS-LRR RGA homologs and CRRSP, whereby pathotype specific effector(s) might target a decoy of RGA homologs, a unique CRRSP, or possibly both, supporting the traditional R-gene guard-decoy hypothesis. Further research is needed to characterize which haplotype 1 gene(s) are truly responsible for ‘Gasaway’ EFB resistance and whether other EFB resistance sources are derived from this same hypothesized molecular mechanism, with R-gene homologs acting in congruence with unique CRRSP proteins. The hazelnut breeding program at OSU has used many different sources of EFB resistance and sequenced their genomes in an effort to expand knowledge of the allelic diversity of putative resistance gene candidates. Future work in determining EFB resistance mechanisms of other *C. avellana* cultivars should be based on comparisons between the pool of R-genes and CRRSP proteins derived from haplotype 1 of ‘Jefferson’ to prospective EFB resistance genes in order to narrow the list of putative candidate genes.

## Conclusions

Here, we report the first haplotype-resolved chromosome-level genome assembly and annotation of the diploid *C. avellana* ‘Jefferson’. BUSCO analysis showed that the genome assemblies and structural annotations were of high quality. The ability of haplotype-phasing to identify parental genic contributions was successfully demonstrated by the complete separation of SI-alleles to their respective parental haplotypes. Furthermore, the region associated with ‘Gasaway’ EFB resistance was remapped with high confidence to the resistant parental haplotype, and several new candidate resistance genes were identified. The molecular mechanism behind ‘Gasaway’ resistance remains to be investigated, however, the RGA cluster in congruence with a cysteine-rich secretory protein provides evidence of a guard model hypothesis. The haplotype-resolved ‘Jefferson’ genome assembly and annotation presented here will serve as a powerful resource for hazelnut breeders and plant scientists in the further development of molecular markers for genomics-assisted breeding and facilitate future studies of *Corylus* biology and genetics.

## Data availability

The haplotype genome assemblies and annotations of *C. avellana* ‘Jefferson’ presented here is available at the United States Department of Energy’s Joint Genomics Institute Phytozome web browser (accepted, pending release) (available The ‘Jefferson’ genome assembly, annotation, and respective read tracks will also be available soon as a genome browser via JBrowse2 at Hazelnutgenomes.oregonstate.edu.

## Acknowledgements

SCT performed flow cytometry, genome assembly, quality assessments, Hi-C guided assembly, synteny analysis, structural and functional annotation, and related analyses of potential candidate genes for resistance and self-incompatibility. JC assisted with Hi-C assembly. JWS conducted remapping of linkage groups and provided insight into candidate gene analysis. KJV coordinated research and provided conceptual guidance, and assisted with assembly, annotation, repeat content, and resistance gene analysis. SAM conceived the study and provided overarching guidance. SCT and KJV authored the manuscript. Oregon State University Center for Quantitative Life Sciences provided the support for computational resources used. All authors approved the final manuscript. The Oregon Hazelnut Commission supported this research.

## Supplemental

**Table S1.** Summary of sequencing data from Illumina, Hi-C, and PacBio platforms.

| Sequencing platform | Sample | Insert length | Sequencing model | Number of reads | Total nucleotides |
| --- | --- | --- | --- | --- | --- |
| PacBio Sequel IIe | 'Jefferson' | >20kb | 2x 8M SMRT cell | 3.64 M | 56.8 Gb |
| Hiseq 4000 | 'OSU 252.146' | 300bp | 2x150 | 295.9 M | 44.38 Gb |
| Hiseq 4000 | 'OSU 414.062' | 300bp | 2x150 | 218.2 M | 32.73 Gb |
| Dovetail Hi-C on Hiseq 4000 | 'Jefferson' | 300bp | 2x150 | 428.46 M | 64.69 Gb |

**Table S2.**
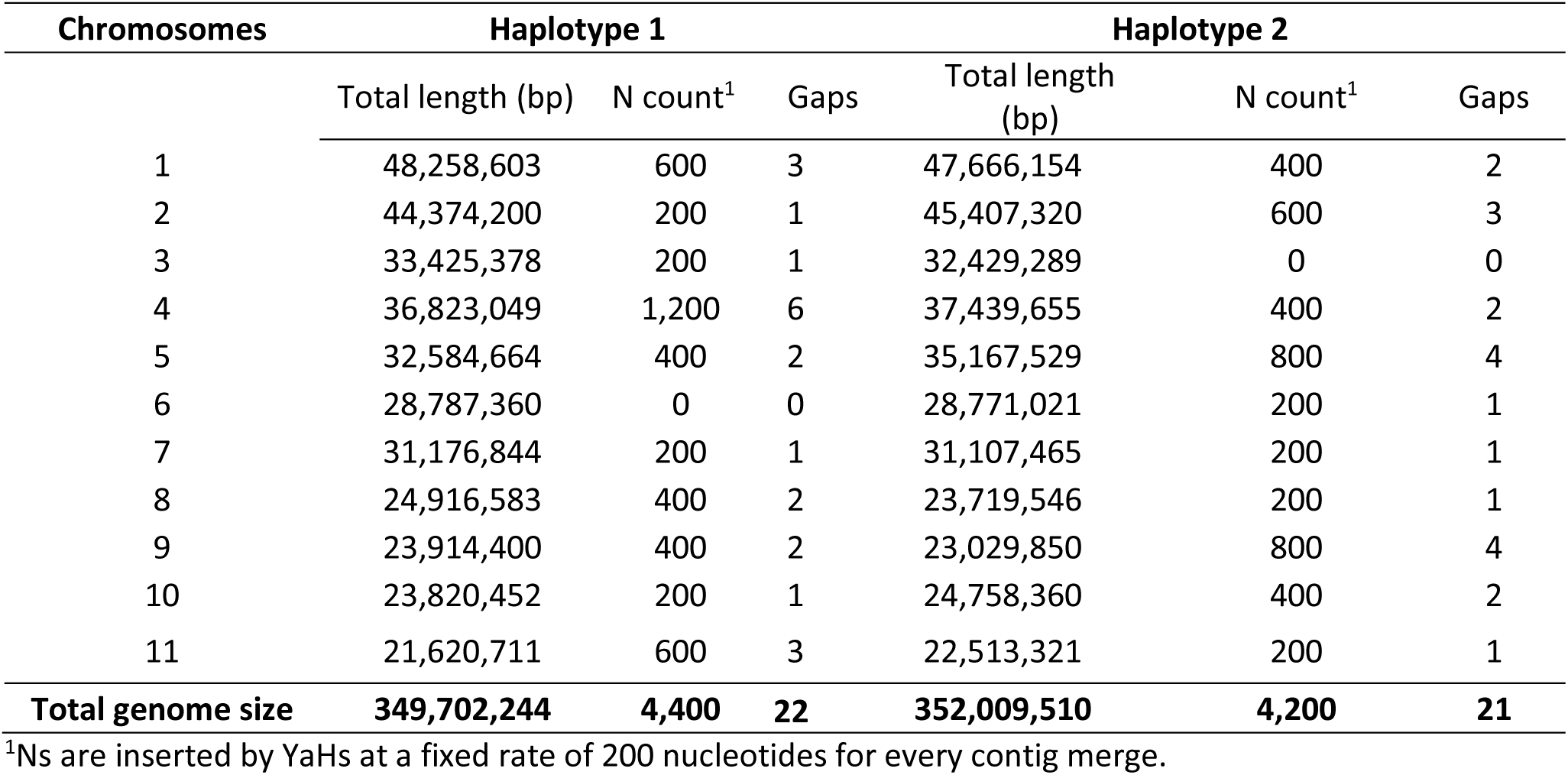
Summary statistics for the eleven haplotype scaffolds corresponding to the ‘Jefferson’ European hazelnut (*C. avellana*) base chromosomes.

**Table S3.** Number and percentage of aligned Illumina 150 bp PE reads derived from ‘Jefferson’ parents to ‘Jefferson’ chromosome-level haplotype-resolved assemblies.

|  | Number and percentage of aligned reads for each parent |  |
| --- | --- | --- |
|  | 'OSU 252.146' | 'OSU 414.062' |
| 'Jefferson' V4 Haplotype 1 | 267,993,778 (90.57%) | 200,943,531 (92.08%) |
| 'Jefferson' V4 Haplotype 2 | 272,354,026 (92.04%) | 199,451,255 (91.39%) |

**Table S4.** Structural variations by SyRI^1^ of ‘Jefferson’ haplotype 1 and haplotype 2.

| Structural variation | Count | Haplotype 1 length (bp) | Haplotype 2 length (bp) |
| --- | --- | --- | --- |
| Syntenic regions | 253 | 277,416,244 | 276,792,668 |
| Inversions | 37 | 6,741,817 | 6,220,076 |
| Translocations | 216 | 41,832,328 | 42,723,238 |
| Duplications (reference) | 259 | 4,088,393 | ----- |
| Duplications (query) | 353 | ---- | 2,863,543 |
| Not aligned (reference) | 593 | 23,471,255 | ---- |
| Not aligned (query) | 677 | ---- | 23,296,270 |

<sup>1</sup>Count table output from default SyRI run, derived from a minimap2 .bam alignment between both haplotypes with the parameter: --eqx.

**Table S5.** Sequence variations by SyRI^1^ of ‘Jefferson’ haplotype 1 and haplotype 2.

| Sequence variation | Count | Haplotype 1 length (bp) | Haplotype 2 length (bp) |
| --- | --- | --- | --- |
| SNPs | 1,593,404 | 1,593,404 | 1,593,404 |
| Insertions | 213,321 | ---- | 2,792,929 |
| Deletions | 150,213 | 2,897,238 | ---- |
| Copygains | 91 | ----- | 471,299 |
| Copylosses | 80 | 253,126 | ----- |
| Highly diverged | 13,753 | 116,506,195 | 116,208,998 |
| Tandem repeats | 22 | 86,034 | 71,726 |

<sup>1</sup>Count table output from default SyRI run, derived from a minimap2 .bam alignment between both haplotypes with the parameter: --eqx.

**Table S6.** ‘Jefferson’ haplotype 1 assembly EDTA^1^ output.

| Class | Number of elements | Length (bp) | Percentage of genome |
| --- | --- | --- | --- |
| **LTR** | **103,937** | **62,391,913** | **17.84%** |
| Copia | 23,301 | 15,899,955 | 4.55% |
| Gypsy | 25,625 | 21,135,697 | 6.04% |
| unknown | 55,011 | 25,356,261 | 7.25% |
| **TIR** | **158,209** | **40,419,512** | **11.55%** |
| CACTA | 33,740 | 9,654,395 | 2.76% |
| Mutator | 77,114 | 17,487,304 | 5.00% |
| PIF_Harbinger | 22,873 | 5,671,550 | 1.62% |
| Tc1_Mariner | 3,700 | 878,845 | 0.25% |
| hAT | 20,782 | 6,727,418 | 1.92% |
| **nonLTR** | **1,065** | **361,136** | **0.10%** |
| LINE_element | 1,031 | 352,462 | 0.10% |
| unknown | 34 | 8,674 | 0.00% |
| **nonTIR** | -- | -- | -- |
| helitron | **35,911** | **9,497,804** | **2.72%** |
| **repeat_region** | **81,361** | **21,122,389** | **6.04%** |
| **Total Genome Masked** | **380,483** | **133,792,754** | **38.26%** |

<sup>1</sup>EDTA run with parameters: --cds --bed --sensitive 1 --analysis 1; the CDS and bed file provide is derived from the BRAKER1/BRAKER2 gene set produced for the respective haplotype.

**Table S7.** ‘Jefferson’ haplotype 2 assembly EDTA^1^ output.

| Class | Number of elements | Length (bp) | Percentage of genome |
| --- | --- | --- | --- |
| **LTR** | **126,989** | **67,657,376** | **19.22%** |
| Copia | 27,892 | 17,200,312 | 4.89% |
| Gypsy | 26,884 | 21,861,582 | 6.21% |
| unknown | 72,213 | 28,595,482 | 8.12% |
| **TIR** | **145,984** | **37,854,874** | **10.75%** |
| CACTA | 27,535 | 7,250,762 | 2.06% |
| Mutator | 74,679 | 18,295,776 | 5.20% |
| PIF_Harbinger | 17,714 | 4,238,613 | 1.20% |
| Tc1_Mariner | 3,122 | 681,523 | 0.19% |
| hAT | 22,934 | 7,388,200 | 2.10% |
| **nonLTR** | **1,100** | **442,005** | **0.12%** |
| LINE_element | 1,047 | 425,994 | 0.12% |
| unknown | 53 | 16,011 | 0.00% |
| **nonTIR** | -- | -- | -- |
| helitron | **39,521** | **8,684,384** | **2.47%** |
| **repeat_region** | **37,689** | **9,592,939** | **2.73%** |
| **Total Genome Masked** | **380,483** | **124,231,578** | **35.29%** |

<sup>1</sup>EDTA run with parameters: --cds --bed --sensitive 1 --analysis 1; the CDS and bed file provide is derived from the BRAKER1/BRAKER2 gene set produced for the respective haplotype.

**Table S8.** Distribution of resistance-like transcripts identified by DRAGO2 among 11 pseudo-chromosomal scaffolds of the ‘Jefferson’ haplotype 1 and haplotype 2 assemblies.

| 'Jefferson'<br>Pseudo-chromosomal<br>scaffolds | CN<sup>1</sup> | CNL<sup>1</sup> | NL<sup>1</sup> | RLK<sup>2</sup> | RLP<sup>2</sup> | TN<sup>3</sup> | TNL<sup>3</sup> | Other<sup>4</sup> | Total |
| --- | --- | --- | --- | --- | --- | --- | --- | --- | --- |
|  | H1/H2 | H1/H2 | H1/H2 | H1/H2 | H1/H2 | H1/H2 | H1/H2 | H1/H2 | H1/H2 |
| 1 | 17/16 | 63/60 | 43/47 | 20/17 | 29/32 | 0/2 | 8/5 | 304/276 | 484/455 |
| 2 | 18/23 | 12/14 | 22/28 | 84/65 | 91/58 | 3/1 | 0/0 | 383/403 | 613/592 |
| 3 | 4/5 | 5/5 | 8/7 | 36/36 | 28/25 | 3/5 | 14/11 | 162/169 | 260/263 |
| 4 | 3/10 | 14/8 | 1/4 | 50/57 | 24/24 | 0/0 | 1/1 | 189/207 | 284/315 |
| 5 | 3/1 | 4/0 | 12/11 | 41/40 | 43/23 | 5/2 | 6/6 | 224/225 | 342/298 |
| 6 | 1/2 | 2/1 | 4/5 | 45/49 | 15/31 | 15/13 | 21/15 | 233/240 | 335/356 |
| 7 | 0/0 | 5/5 | 2/1 | 66/71 | 44/62 | 0/0 | 0/0 | 165/179 | 284/320 |
| 8 | 5/8 | 11/20 | 5/5 | 39/37 | 70/58 | 14/10 | 47/31 | 166/169 | 360/345 |
| 9 | 11/14 | 10/9 | 3/6 | 15/21 | 16/21 | 1/1 | 12/10 | 126/128 | 196/211 |
| 10 | 2/1 | 11/9 | 6/2 | 45/50 | 63/66 | 0/0 | 2/3 | 136/147 | 265/280 |
| 11 | 0/0 | 2/2 | 0/0 | 45/45 | 22/23 | 0/0 | 0/0 | 127/154 | 197/224 |
| **Total** | **66/80** | **139/133** | **117/122** | **486/445** | **445/**<br>**423** | **41/34** | **111/82** | **2,215/**<br>**2,297** | **3,620/**<br>**3,659** |

<sup>1</sup>Coiled-coil nucleotide binding site [(CC-NBS (CN))]; CC-NBS-leucine rich repeat [(CC-NBS-LRR (CNL))]; NBS-LRR (NL).
<sup>2</sup>Receptor-like Kinase (RLK); Receptor-like Protein (RLP).
<sup>3</sup>TIR-NBS (TN); TIR-NBS-LRR (TNL).
<sup>4</sup>Includes kinases (K), NBS (N), LRRs (L), CKs, CTs, CTLs, Lysine motif containing proteins (LYK and LYP) and Lectin-like motif containing proteins (LECM).

**Figure S1.**
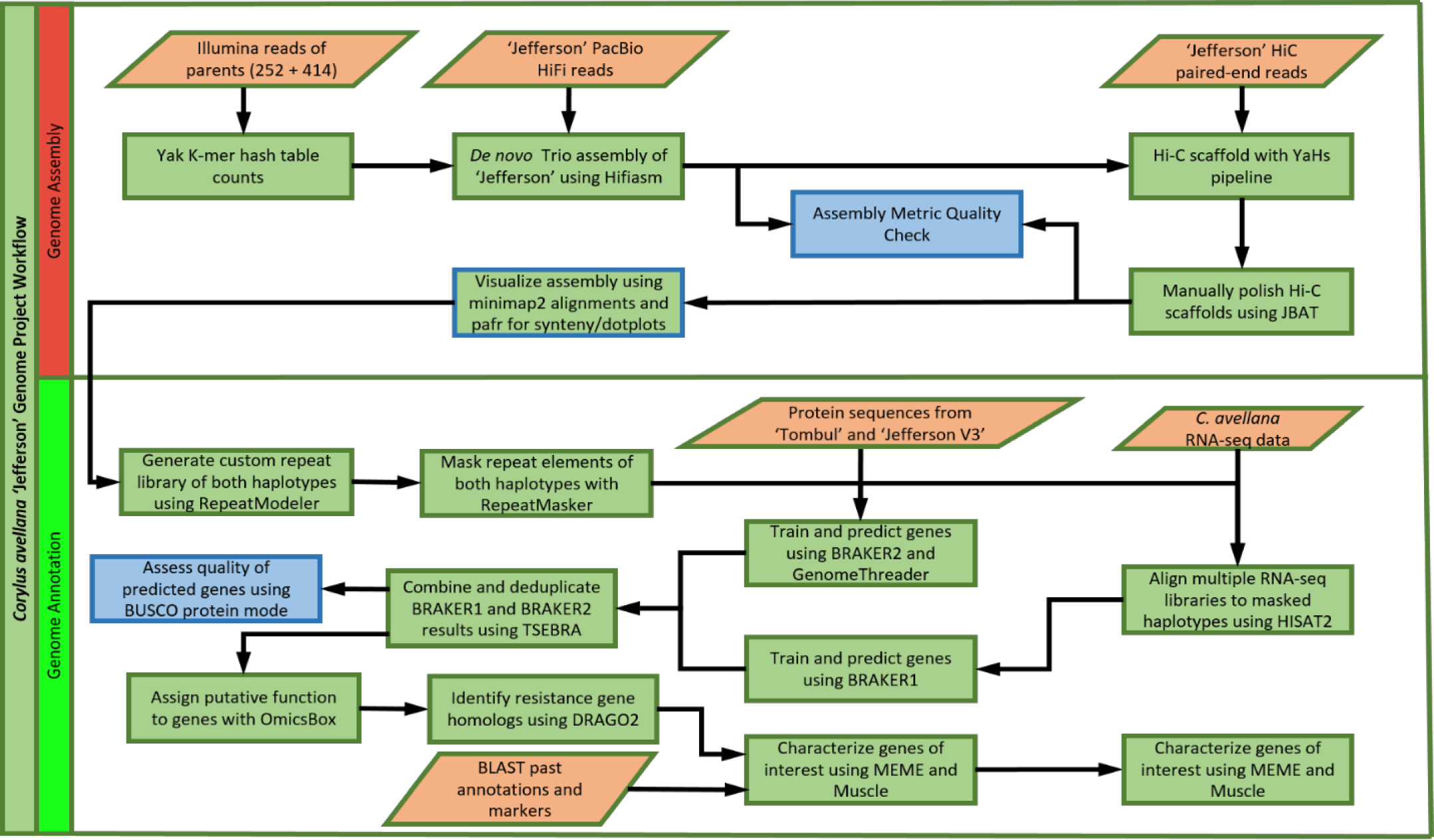
Genome assembly and annotation workflow of *C. avellana* ‘Jefferson’. Figure shows the genome assembly and annotation pipeline with processes shown in green, extraneous data in orange and quality checks in blue.

**Figure S2.**
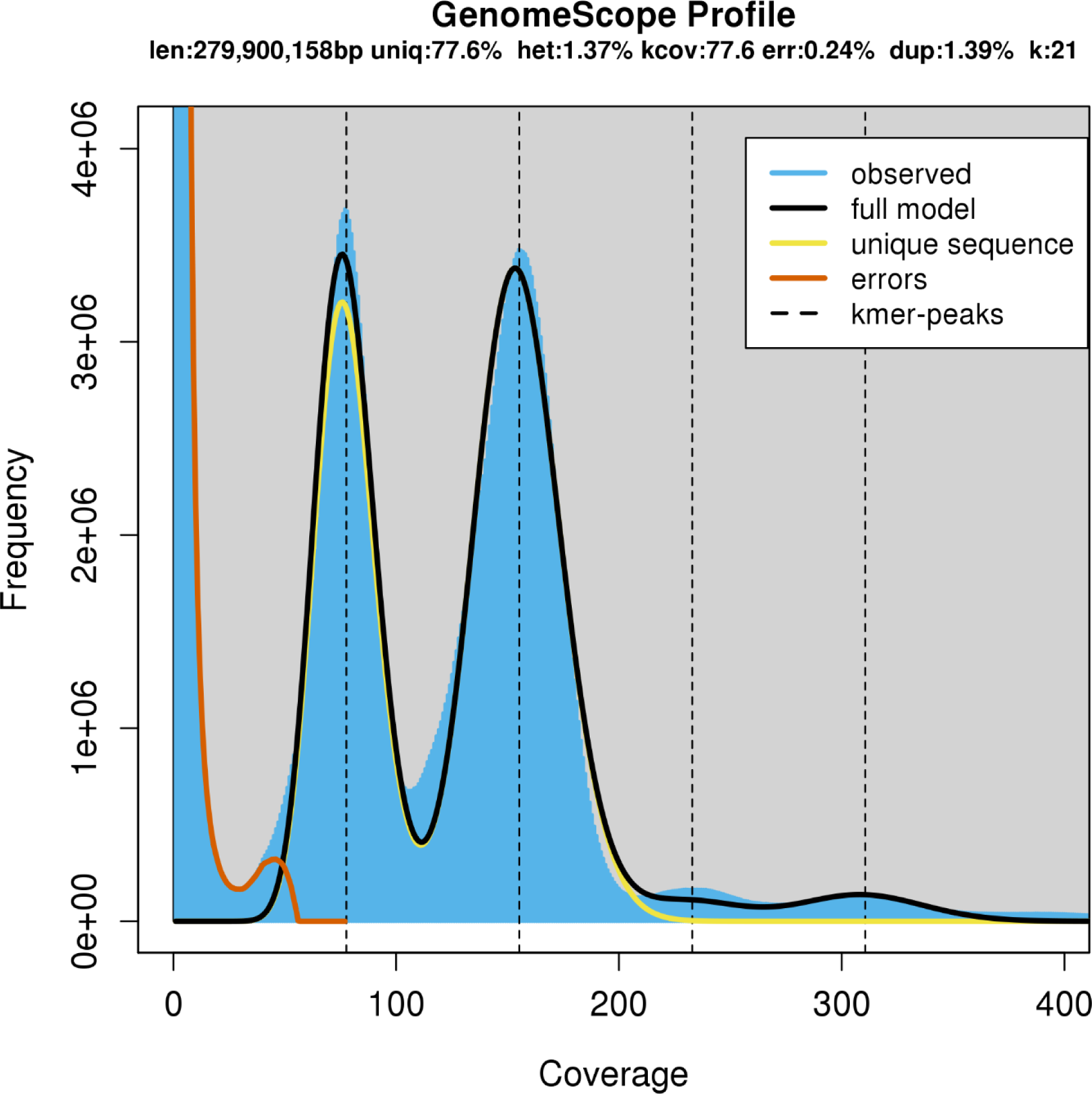
GenomeScope result of raw ‘Jefferson’ PacBio HiFi reads for k-mer length = 21. GenomeScope output derived from jellyfish count -C -m 21 -s 1000000000 and jellyfish histo.

**Figure S3.**
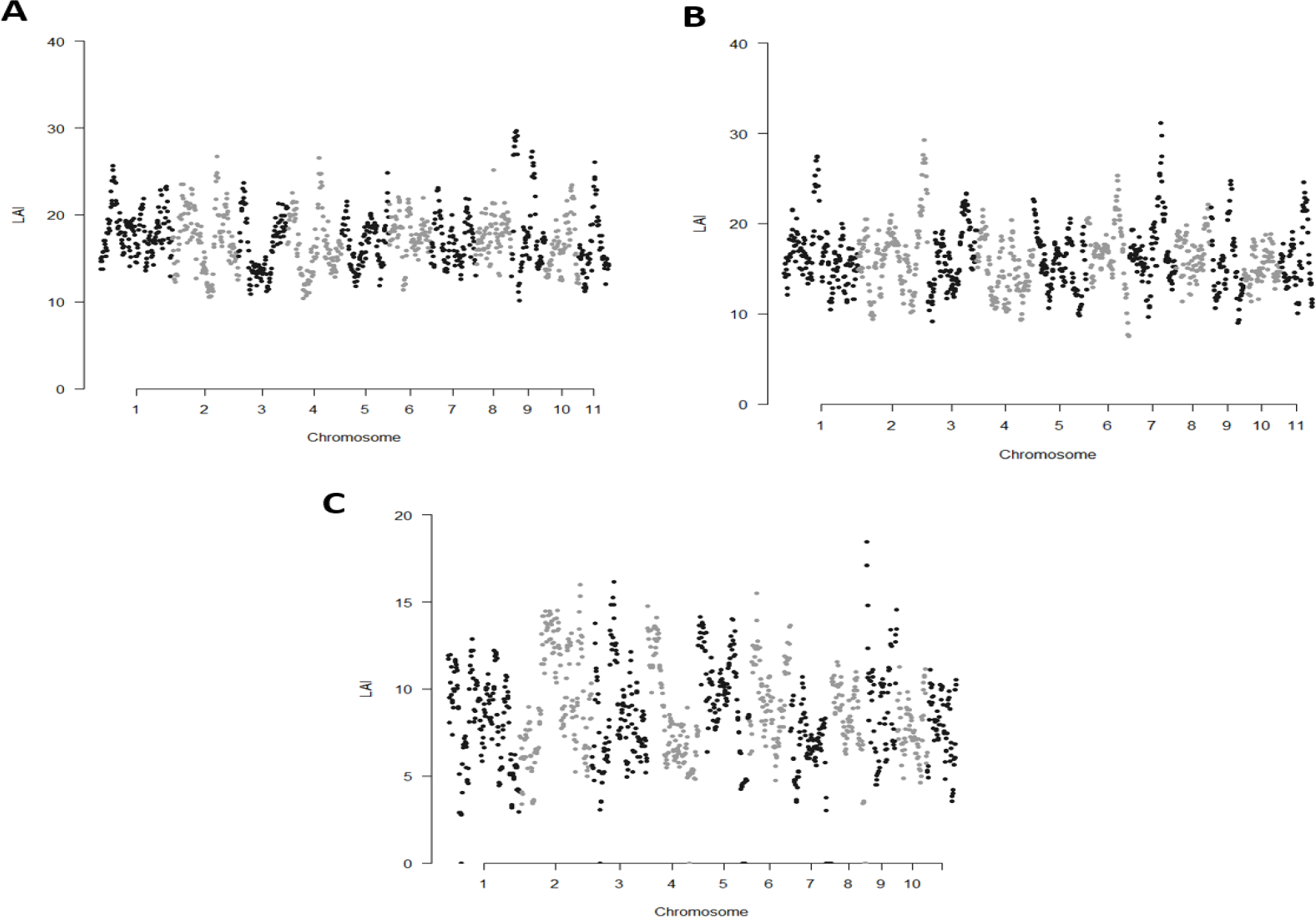
LAI scores of *C. avellana* ‘Jefferson’ haplotypes and ‘Tombul’. LAI scores were obtained by LTR_Retriever from a concatenated set of LTRs derived from LTR harvest and LTR_FINDER_parallel for each respective assembly. Each dot represents LAI score of a 3 Mb-sliding window with 300-Kb increment, adjusted by the whole-genome LTR identity. (**A**) ‘Jefferson’ haplotype 1 genome assembly. (**B**) ‘Jefferson’ haplotype 2 genome assembly. (**C**) ‘Tombul’ genome assembly.

**Figure S4.**
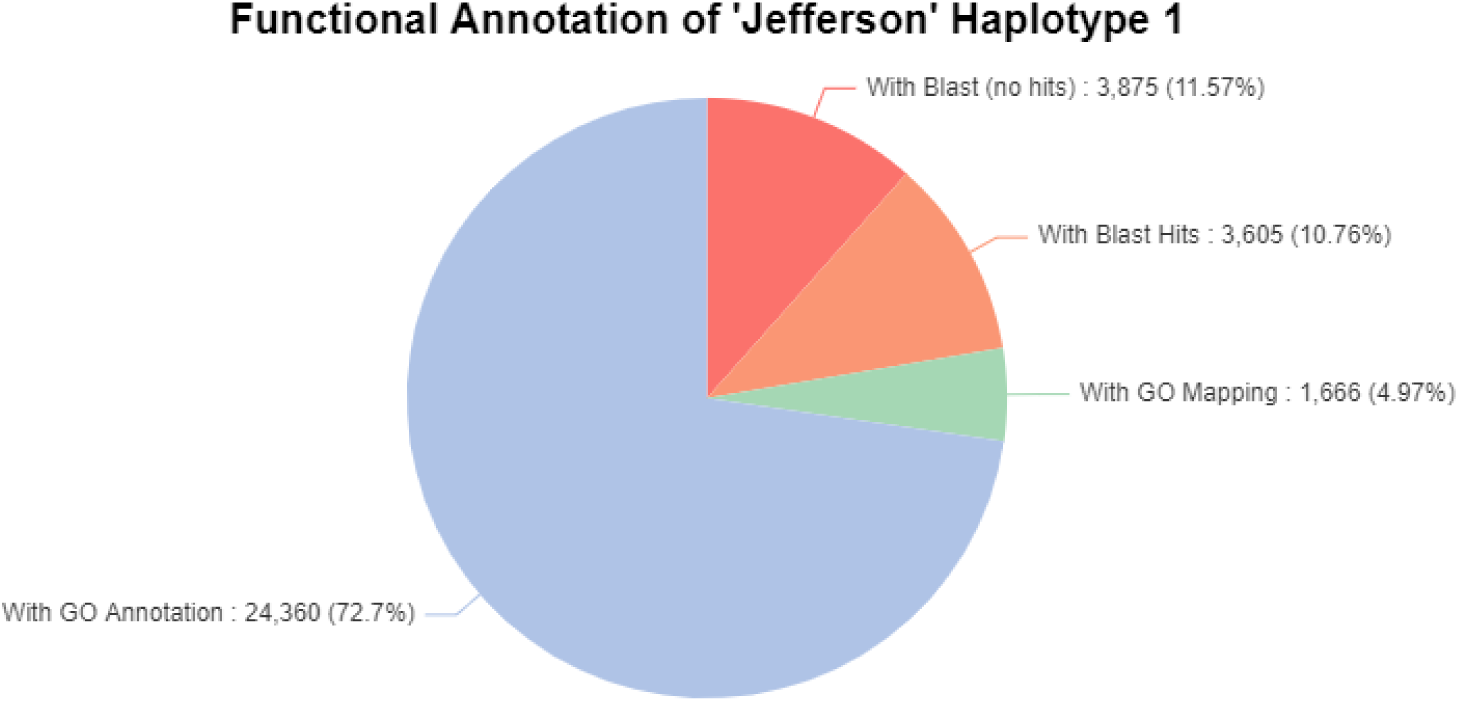
OmicsBox summary metrics of ‘Jefferson’ haplotype 1 functional annotation. Pie chart shows total distribution of OmicsBox functional annotation performed on haplotype 1 amino acid transcripts of ‘Jefferson’. In red are transcripts that received no BLAST hits from the database and thus have unknown function; orange are transcripts that received only BLAST hits; green are transcripts that had GO terms associated with the initial BLAST database search; blue is transcripts that received GO annotation descriptions.

**Figure S5.**
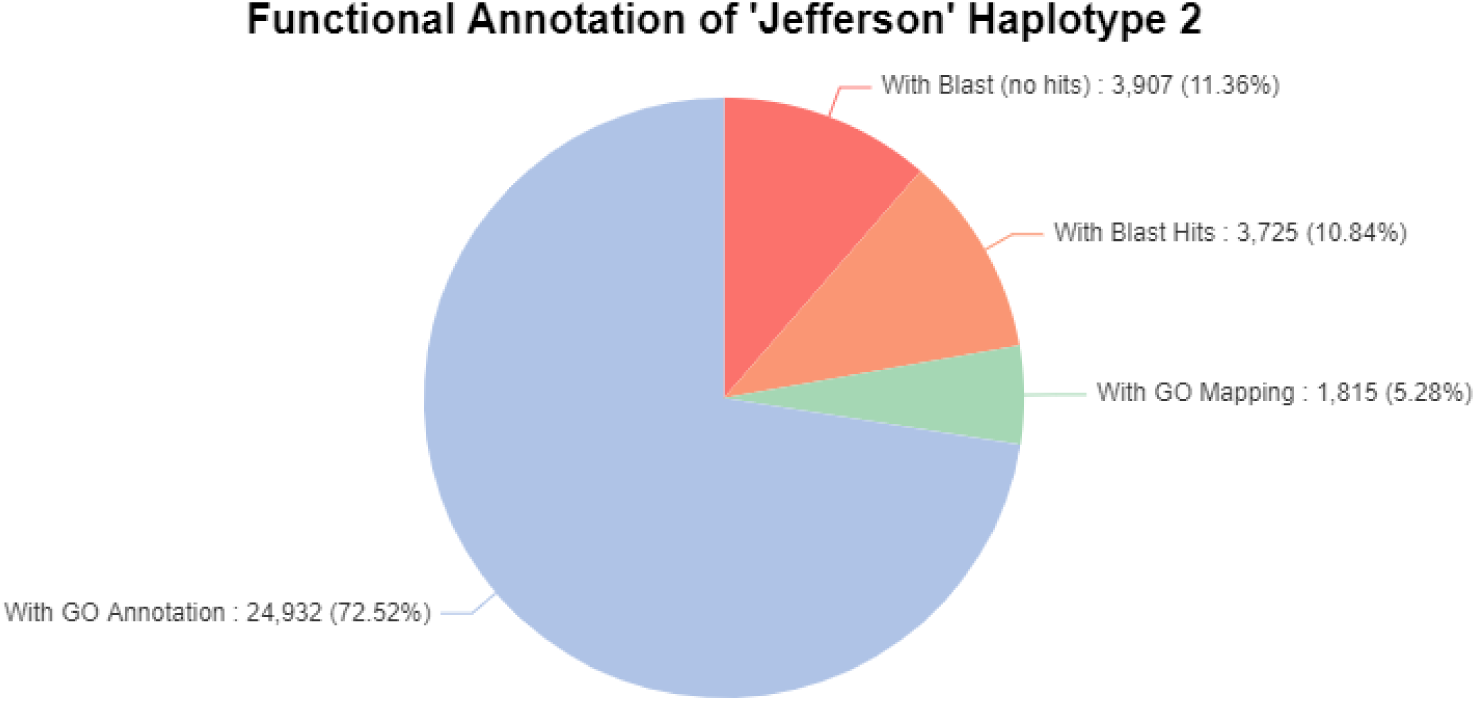
OmicsBox summary metrics of ‘Jefferson’ haplotype 2 functional annotation. Pie chart shows total distribution of OmicsBox functional annotation performed on haplotype 2 amino acid transcripts of ‘Jefferson’.

